# Directional branch migration remodels the meiotic Holliday junction landscape

**DOI:** 10.64898/2026.04.08.717233

**Authors:** Martin Xaver, Anura Shodhan, Nataliia Kashko, Michael Lichten, Joao Matos

**Affiliations:** Max Perutz Labs, Vienna BioCenter, A-1030 Vienna, Austria; University of Vienna, Vienna, Austria; Laboratory of Biochemistry and Molecular Biology, Center for Cancer Research, National Cancer Institute, NIH, Bethesda, MD 20892, USA; Vienna BioCenter PhD Program, a Doctoral School of the University of Vienna and the Medical University of Vienna, A-1030 Vienna, Austria

**Keywords:** Holliday junction pulldown, HJSeq, Holliday junction profiles, budding yeast, meiosis, pachytene, homologous recombination, crossover, ZMM proteins, Bloom’s helicase, DNA repair

## Abstract

Holliday junctions (HJs) are four-way branched DNA intermediates in double strand break (DSB) repair by homologous recombination. During meiosis, HJs are resolved predominantly as crossovers that are necessary for correct chromosome segregation. However, the genome-wide behavior of meiotic HJs has remained inaccessible without direct mapping. By a HJ-binding peptide approach (HJSeq), we capture the genome-wide distribution of these crucial intermediates for the first time. We show that during pachytene, the stage before crossover formation, HJs undergo sustained directional branch migration, reorganizing the HJ-landscape in a transcription-linked manner, away from DSB sites towards convergent transcription sites. The presented findings establish pachytene as an active remodeling phase in which HJ branch-migration accommodates an active chromatin environment to support regulated crossover formation and faithful chromosome segregation.

**Highlights:**

1. HJSeq enables genome-wide mapping of meiotic Holliday junctions
2. HJ formation propensity is determined by DSB hotspot strength
3. Meiotic HJs undergo long-range, transcription-linked branch migration
4. Directional HJ migration away from DSB sites is facilitated by the ZMM pathway

## INTRODUCTION

Holliday junctions (HJs) are four-way intermediates in homologous recombination that connect recombining partner molecules by a reciprocal swap of base pairing strands ^1^. HJs connecting homologs (homologous chromosomes of different parental origin) have an essential function in meiosis, where their nucleolytic resolution as crossovers ensures proper homolog segregation ^2^. In contrast, unresolved or incorrectly placed HJs endanger genomic integrity ^3^. Thus, while somatic cells utilize helicases and endonucleolytic resolvases to prevent HJ accumulation ^3^, meiotic cells facilitate the stabilization of HJs and their crossover specific resolution using the conserved ZMM pathway (named for the budding yeast component proteins Zip1-4, Spo16, Mer3, and Msh4/5) ^4^.

This has been best characterized in budding yeast, where recombination is initiated in meiotic prophase I by about 200 DNA double strand breaks (DSBs), formed by the Spo11 complex at so called hotspots that have a punctuate distribution across the genome ^5,6^. While all DSBs are repaired by homologous recombination, a fraction is transformed by the ZMM pathway into crossover-designated intermediates that contain two HJs (double Holliday junctions, dHJs, Figure 1A) ^4^. The ZMM pathway ensures that crossovers occur at least once per homolog pair, and with a sparse and even distribution via a phenomenon termed interference ^2^.

**Figure 1.**
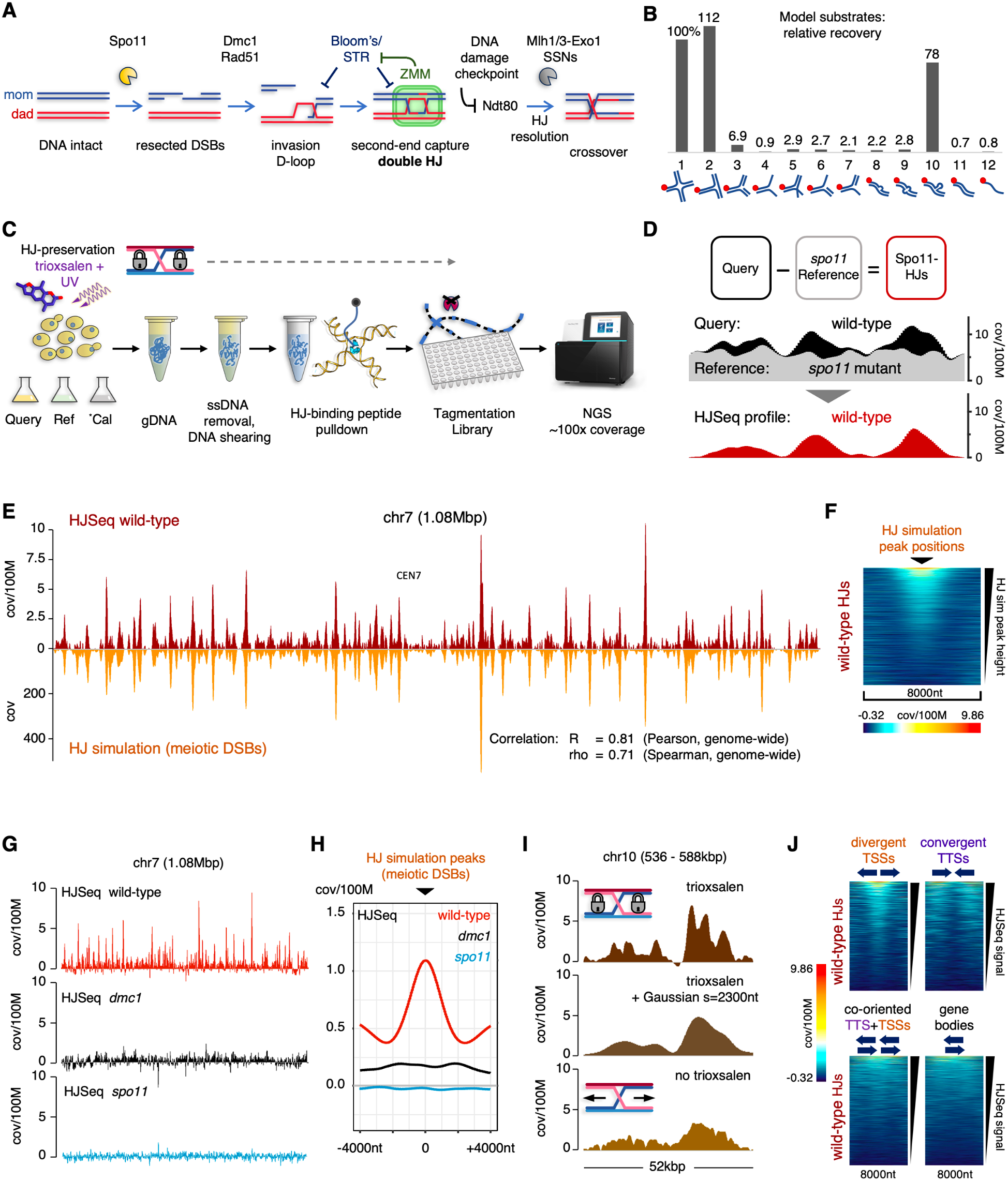
HJ formation propensity follows DSB hotspot strength. (A) Simplified meiotic crossover recombination pathway: Spo11 forms meiotic DNA double strand breaks at hotspots, Dmc1 (and Rad51) catalyze invasion of repair template DNA in homologous recombination, ZMM pathway protects some intermediates against Bloom’s complex (STR) to form crossover-designated double Holliday junctions. DNA damage checkpoint release activates Ndt80 transcription factor expression, committing to HJ resolution into crossovers (Mlh1/3-Exo1 ZMM HJ resolution complex, SSNs structure selective backup nucleases). (B) HJ binding peptide-based pulldown performance on model substrates (2pmol Cy5-labelled model over 3.4µg dsDNA): 1) Branch-migrating HJ, 2) closed 3-way junction, 3) full double stranded fork, 4) flayed fork, 5) double flap junction, 6) 3’ flap DNA, 7) 5’ flap DNA, 8) tetraloop hairpin, 9) 4nt bubble on tetraloop, 10) mini-stemloop, 11) double stranded DNA, 12) single stranded DNA (see Figure S1C for details). (C) Complete HJSeq workflow for budding yeast meiosis: Synchronous meiotic cultures of SK1 *S.cerevisiae*: Query and Reference (*spo11*) strains, optionally IFO1815 *S.mikatae* for calibration, are trioxsalen-treated to form interstrand DNA crosslinks that lock HJs in place. Genomic DNA is isolated under HJ-preserving conditions (Allers and Lichten 2000) with *S.mikatae* calibration spike-in added following the occupancy ratio logic of Hu et al. 2015. DNA is randomly sheared to a median fragment size by weight of ∼750bp, and ssDNA and RNA digestion (Mung Bean Nuclease) reduces sample complexity. HJ DNA is enriched by utilizing the HJ binding peptide-based pulldown (see B). Libraries are prepared by Tn5 tagmentation for sub-fragments devoid of trioxsalen from input and eluate. Paired-end sequencing of Query and Reference Input (∼25x coverage) and Eluate (∼100x coverage) is performed. (D) Query and Reference Eluate coverages are scaled based on the calibration spike-in (cal), or *spo11* reference via Signal Extraction Scaling (SES). Subtraction of scaled Eluate Reference from Query yields differential Spo11-dependent HJ profiles, which are denoised and baseline adjusted based on recombination coldest regions. A region of chr15 (460-480kbp) in wild-type 4h (cal) is shown. Y-axis: calibrated nucleotide coverage per 100M. (E) Chromosome 7 (1.08Mbp), wild-type 4+4.5h (averaged, calibrated) HJSeq (top, dark red, 150nt binning, denoised) compared to DSB-based HJ simulation (bottom, orange, Gaussian smoothed [s=1000nt] Spo11-oligo data SRR7811345 from Murakami et al. 2020) with y axis scale normalized to total wild-type HJSeq signal. Single nucleotide coverage is reported. (F) Genome-wide pile-up heatmap for wild-type 4+4.5h HJSeq (averaged, calibrated) coverages at HJ-simulation peak positions (n=2122). Linear color-to-coverage relation as indicated, scaled to full displayed coverage range. Pile-up heatmap coverages have lower cutoff applied. Coverage: single nucleotide coverage per 100M. (G) Whole chromosome 7 HJSeq profiles, calibrated. wild-type 4h (red), *dmc1* 5+6h (merged, averaged) (black), *spo11* 4.5h (turquoise). Coverage: single nucleotide coverage per 100M. (H) Genome wide mean coverage pile-up analysis relative to HJ-simulation peak positions. Same data as in (G). Coverage: single nucleotide coverage per 100M (calibrated). (I) HJSeq signals spread out in absence of trioxsalen interstrand crosslinks. Section of chromosome 10 (52kbp). Top: Standard HJSeq (trioxsalen treated). Middle: Top, but Gaussian smoothed s=2300nt to best Pearson correlation with Bottom (see Figure S2G). Bottom: HJSeq performed without trioxsalen treatment. Samples were obtained and treated in parallel throughout (HJSeq *ndt80* 6h, calibrated). (J) Genome wide pile-up heatmaps for HJSeq wild-type 4+4.5h coverages, for indicated intergenic regions (mid points) as well as gene body positions. Divergent TSSs: n=1435. Convergent TTSs: n=1430. Co-oriented TTS+TSSs intergenic regions: n=2644. Gene bodies: n=5505. Linear color-to-coverage scale as in (F). X-axis mid-point of heatmap is mid-point position of intergenic regions and gene bodies. Regions (y-axis) sorted by total HJSeq signal for all heatmaps. Pile-up heatmap coverages have lower cutoff applied. Coverage: single nucleotide coverage per 100M.

HJs persist for a considerable time during the pachytene stage of meiotic prophase I, during which chromosomes are organized as axis-loop domains, with homolog axes tightly paired along their length (synapsed) by a proteinaceous structure called the synaptonemal complex (SC), whose establishment and maintenance requires crossover-designated intermediates formed in the ZMM pathway ^2,7,8^. Pachytene lasts about one to two hours in budding yeast, many hours in plants, and several days in mammals ^9-11^. During this period, recombination intermediates are gradually reduced to the essential minimum of crossover-precursors ^12^. The underlying dynamics of these processes of HJ maturation, selection and turnover remain poorly understood, largely because of the complexity of events and pathways involved, the rare and transient nature of these intermediates, and a lack of tools to directly detect and measure HJs at high throughput and resolution.

To address this latter deficit, we developed a novel technique (HJSeq) that uses a class of HJ-binding peptides (Figure S1A) ^13,14^ to enrich HJ intermediates from bulk genomic DNA, followed by high-throughput sequencing and computational mapping. Using budding yeast meiosis, we obtained the first genome-wide maps of meiotic HJs in any organism, enabling direct and quantitative comparison of HJ landscapes in wild type and mutants. The analysis reveals extensive branch migration of HJs on pachytene chromatin in directions that mirror the transcriptional landscape, leading to a pachytene duration dependent, highly stereotypical spatial reorganization of recombination intermediates. Disrupting normal meiotic HJ metabolism prevents this global redistribution, suggesting an integral role for directional branch migration in meiotic recombination.

## RESULTS

### Local Holliday junction abundance follows DSB hotspot strength

Genome-wide levels and distribution of meiotic HJs have remained elusive. We therefore developed HJSeq to directly capture HJs genome-wide. In HJSeq (outlined in Figure 1C), genomic DNA is isolated using conditions (trioxsalen DNA interstrand crosslinking and CTAB extraction) that preserve HJs by suppressing spontaneous branch migration ^15,16^. Sources are synchronous meiotic cultures of both a query and a DSB-deficient (*spo11*) reference strain; for calibrated experiments ^17^, a common spike-in sample from meiotic *Saccharomyces mikatae* is included in all samples. The DNA is enzymatically sheared to a median of ∼750bp, and ssDNA and RNA are removed by Mung bean nuclease digestion followed by ultrafiltration (Figure S1D, and S1E). HJ-containing DNA is enriched by pull-down and release with a biotin-linked HJ-binding peptide (Figure 1B, S1B, and S1C). Tn5-mediated tagmentation is used to generate libraries with sequenceable subfragments devoid of trioxsalen adducts (Figure S1F), and paired-ended sequencing is performed to about 100-fold coverage. Coverages for query and reference samples are scaled to a normalized nucleotide coverage, using either the *S.mikatae* calibration spike-in (“cal”), or Signal Extraction Scaling (“SES”) for experiments lacking a spike-in ^18^. Since processes like DNA replication and transcription can form structures that mimic recombination-formed HJs, a differential profile of meiosis-specific HJs is obtained by subtracting an appropriately scaled reference (*spo11*) profile from the query profile (Figure 1D), using a zero-coverage baseline determined from the genome regions coldest for meiotic recombination (Figure S1H).

We find that wild-type HJSeq signal patterns closely correspond to those of meiotic DSBs (Figure 1E, 1F, S2A, and S2B) ^6,19,20^. The greatest genome-wide correlation between wild-type HJSeq and DSB profiles (Pearson R=0.81, Spearman rho=0.71) was obtained by Gaussian smoothing of Spo11-oligo data ^21^ with a sigma of ∼1000nt (Figure S1G; hereon referred to as ‘HJ simulation’), which, after accounting for fragment-induced signal broadening, matches previous estimates of DSB-to-HJ distances (Figure S2F) ^20^. Accordingly, both DSB and wild-type HJSeq signals preferentially mapped to divergent transcription start sites (TSSs), which are enriched for strong recombination hotspots, while convergent transcription termination sites (TTSs) were particularly cold (Figure 1J, and S3D) ^6^. Crucially, HJSeq signals were dependent on Spo11, which forms DSBs, and Dmc1, which catalyzes the first step of DSB repair by homologous recombination (Figure 1A, 1G, 1H, and S2C–S2E) ^22-25^. Furthermore, when DNA was prepared without trioxsalen crosslinking, a pronounced peak broadening was observed, approximated by additional Gaussian smoothing of the crosslinked profile by s=∼2300nt (Figure 1I, and S2G), as would be expected from stochastic branch migration of HJs during DNA processing in the absence of crosslinks ^16,26^. Together with genetic analyses and physical Southern blotting assays presented below (Figure S2H, 3A–3C, and Figure 4), these observations indicate that HJSeq specifically detects meiotic HJ intermediates, enabling direct, genome-wide mapping of their local distributions. Thus, the strong genome-wide correlation between wild-type HJSeq and Spo11-DSB data indicates that local HJ-formation propensity is determined primarily via Spo11-DSB hotspot activity.

We note that unsubtracted HJSeq data contained substantial meiotic recombination-independent signal in ∼2.7% of the genome. This signal mapped to palindromic TA(n>10)-runs and tandem repeat sequences with self-folding potential, predominantly in subtelomeric and DSB-cold regions ^6^ (Figure S2I). These regions were excluded from further analysis.

### HJs relocate towards convergent transcription sites during extended pachytene

Under the sporulation conditions used here, wild-type budding yeast of the SK1 background spends 1-2 hours in pachytene, the meiotic stage where HJs are present ^9,15,27,28^. In the culture, peak pachytene occurs within the ∼4-5h window after sporulation induction (Figure S3F–S3N), limiting the HJ age-range that can be resolved. Because pachytene exit and HJ resolution depend on the transcription factor Ndt80 (Figure 1A) ^27,29,30^, *ndt80* mutants provide a tractable way to prolong pachytene and examine the effects of extending HJ lifespan.

We performed HJSeq on synchronized *ndt80* cultures after prolonged pachytene duration (7.5h in sporulation medium, SPM; biological replicate 7.0h). As expected for a culture mostly composed of pachytene cells, HJ levels were elevated. However, the spatial distribution of HJ signal differed extensively from that seen in 4h samples from wild type. At a broad scale (tens of kilobases), the *ndt80* HJ landscape still reflected the underlying regional recombination activity, as defined by wild-type HJSeq and Spo11 DSB distributions. In contrast, locally, HJSeq peaks were displaced away from DSB regions, typically by multiple kilobases (Figure 2A, and S3A). Samples taken at 1h intervals across *ndt80* pachytene (4.5–7.5h) showed a progressive redistribution, with signal gradually leaving hotspot-proximal regions and accumulating at discrete endpoints (Figure 2E). These endpoints reveal a stereotypical and pervasive pattern, colocalization with chromosome axis sites, as marked by the axis protein Hop1, that flank recombination hotspots ^31^ (Figure 2A, 2E, and S3A). These axis sites are recombination cold and generally contain convergent transcription termination sites (TTSs) (Figure 2B) ^6,32-36^.

**Figure 2.**
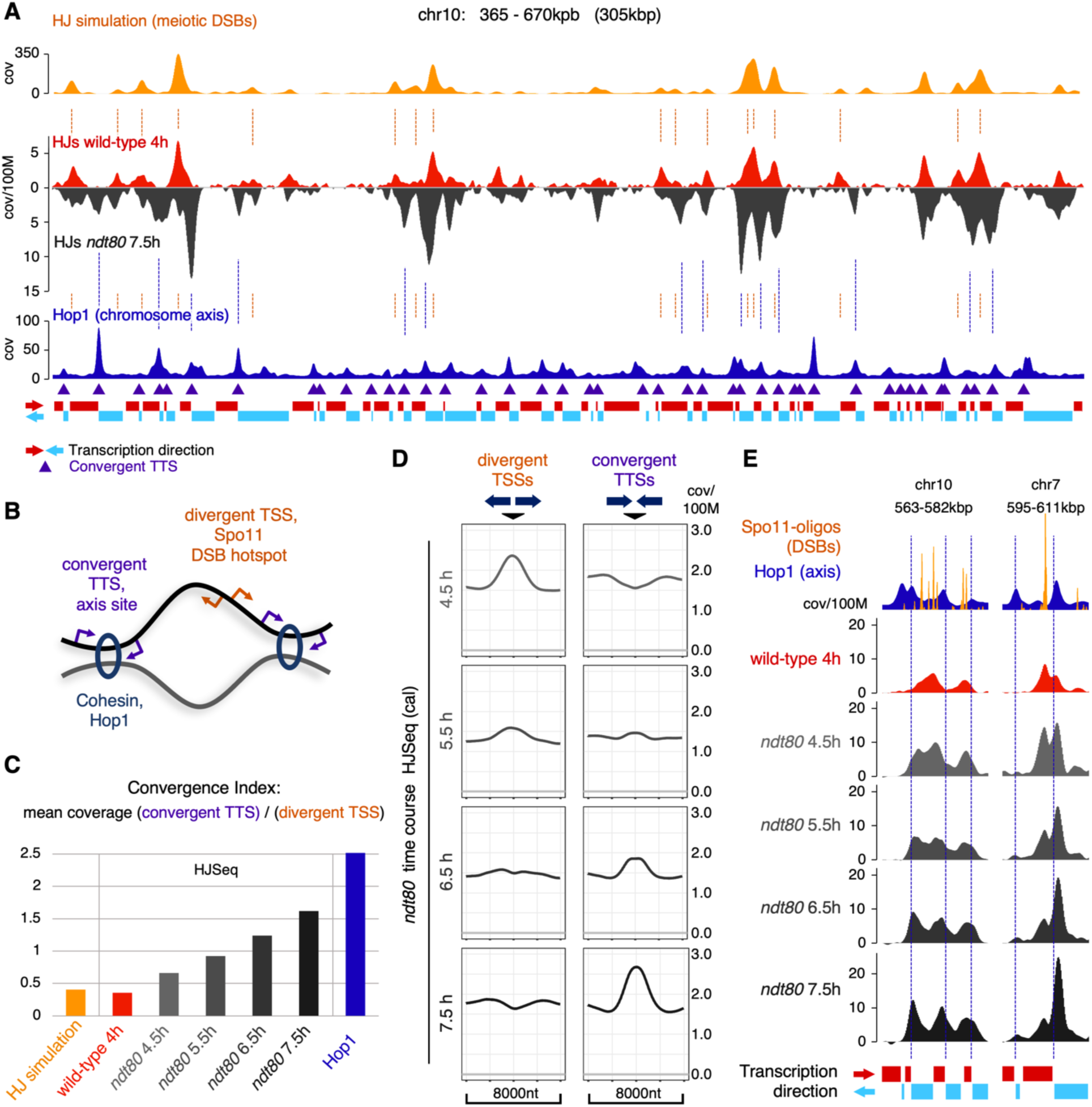
Pachytene HJs relocate towards convergent transcription sites. (A) Right arm of chromosome 10. Orange: HJ simulation (wild-type 4h) derived from Spo11-oligo data. Red: HJSeq wild-type 4h (cal). Dark gray: HJSeq pachytene arrested *ndt80* 7.5h (cal). Blue: Hop1 (wild-type 4h, ChIP-Seq) (Shodhan et al. 2021) indicating chromosome axis sites. Violet triangles: Convergent TTSs. Red and blue rectangles: transcription direction as per underlying ORF and co-oriented ORF runs (red: transcribed right-wards; blue: transcribed left-wards). HJSeq coverage: nucleotide coverage per 100M. (B) Axis-loop domains of meiotic chromosomes are defined by the transcriptional landscape. DSB hotspots are in promoters of DNA loops, particularly divergent TSSs. Axis sites are defined by convergent TTSs carrying meiotic cohesin and axis proteins (including Hop1). Sister chromatid DNA pairs (in black and gray) are tethered by cohesin at chromosome axis sites. Arrows indicate transcription direction (shown for one chromatid only). (C) Convergence Indices for the indicated genome-wide data. Ratio of mean coverage for convergent TTSs (n=1414, 501nt windows) over mean coverage for divergent TSSs (n=1419, 501nt windows). HJ simulation from Spo11-oligo data as above. Hop1 (chromosome axis marker) from ChIP-Seq data as above. (D) Genome-wide HJSeq (cal) *ndt80* time course (4.5-7.5h) pile-up mean coverage data at divergent TSS (n=1419) and convergent TTS (n=1414) positions. Coverage: nucleotide coverage per 100M (average for all regions). (E) Gradual HJ relocation to convergent transcription sites in pachytene for two example regions from calibrated HJSeq *ndt80* time course (4.5-7.5h, gray to black). Orange: Spo11-oligo data wild-type 4h (Murakami et al. 2020). Blue: Hop1 wild-type 4h ChIP-Seq, as above. Red: HJSeq wild-type 4h (cal), as above. Transcription direction as per underlying ORF and co-oriented ORF runs (red: transcribed right-wards; blue: transcribed left-wards). HJSeq coverage: nucleotide coverage per 100M.

Genome-wide inspection also showed that HJ relocation follows the underlying transcriptional landscape. Initially, HJSeq signal is found predominantly at divergent TSSs, which contain the strongest Spo11 DSB hotspots (Figure 1J). The HJSeq signal then steadily redistributes over time towards convergent TTSs (Figure 2D, S3B, and S3C). HJSeq signal originating in intergenic regions between co-oriented genes likewise follows the ORF-pair orientation (Figure S3E).

Overall, prolonging pachytene produces a genome-wide reconfiguration of the HJ landscape, shifting enrichment away from hotspot-proximal regions and towards convergent TTSs in a manner aligned with transcriptional architecture. To quantify this transition, we defined a Convergence Index as the ratio of genome-wide mean coverage at convergent TTSs to mean coverage at divergent TSSs (501nt windows) (Figure 2B, and 2C). This metric increases progressively over time during *ndt80* arrest, readily capturing the global and steady shift towards convergent sites.

### Meiotic Holliday junctions branch migrate directionally in pachytene

HJs can move along DNA by a process called branch migration ^1^. In vivo, however, spontaneous migration is limited by topological constraints ^37-39^. Nonetheless, our HJSeq data indicates that bulk HJ relocation occurs efficiently during the extended pachytene of *ndt80* mutants, on average ∼2kbp and up to 5kbp, possibly limited only by the quasi-regular presence of chromosome axis sites. To substantiate that HJSeq signals seen during extended pachytene indeed are a signal of HJs between recombination partners, we performed Southern blot assays of joint molecules (JMs) at 13 loci (1.9-2.2kbp windows); eight located within regions where HJSeq patterns deviate from expectations based on the Spo11-oligo derived HJ simulation (Figure 3B, and S4A). The JM levels in DNA from 7h *ndt80* cultures showed a strong correlation with respective HJSeq patterns (Figure 3C; coefficient of determination R^2^=0.89, Figure S4B). Similar results were obtained for JMs (12 of above loci assayed) in an independent time course without trioxsalen crosslinking of DNA, albeit JM levels were lower, reflecting reduced HJ stabilization (Figure 3C, and S4B). In wild-type cultures, HJSeq and JM levels displayed similar kinetics with respect to meiotic progression (Figure S3F–S3J). In fact, HJSeq patterns from wild type 4h cultures already displayed signs of directional HJ relocation (Figure 3B, and S4A), consistent with the general movement of HJs away from DSB sites. We considered the possibility that the apparent HJ redistribution seen during extended pachytene might be due to de novo DSB formation. Pachytene-arrested *ndt80* cells continue to make DSBs, albeit at reduced levels compared to early meiotic prophase I ^27,29,40^. A shift towards convergent TTSs, in either DSB formation or stabilization of newly-formed HJs, could account for the observed HJ landscape changes. To test this suggestion, we rapidly shut off DSB formation by rapamycin-mediated “anchor-away” of the Spo11 cofactor Rec104 ^41^, creating a defined HJ population that can be followed as it ages (Figure 3D, and S4C–S4E). We reasoned that, if HJSeq signal redistribution over time reflects new HJ formation, it should not be seen in the absence of new DSB formation. HJSeq signals from DNA collected 30 minutes after DSB shutoff (“young HJs”) presented a HJ population with incipient re-location appearance, but HJSeq signals at divergent TSSs still dominated. In contrast, in DNA collected 3.5h later (“old HJs”), HJSeq distribution patterns displayed pronounced accumulation at convergent TTSs (Figure 3E–G). Importantly, HJSeq signals for “old HJs” accumulated to similar levels at convergent TTSs as for the DSB proficient control (“accrued HJs”), and showed only a modest reduction in overall signal, attributable to a lack of new DSB formation (Figure 3E–3G, S4F, and S4H). Thus, the HJSeq signal that accumulates over time at convergent TTSs cannot be due to new HJ formation, and thus must derive from intermediates that formed elsewhere.

**Figure 3.**
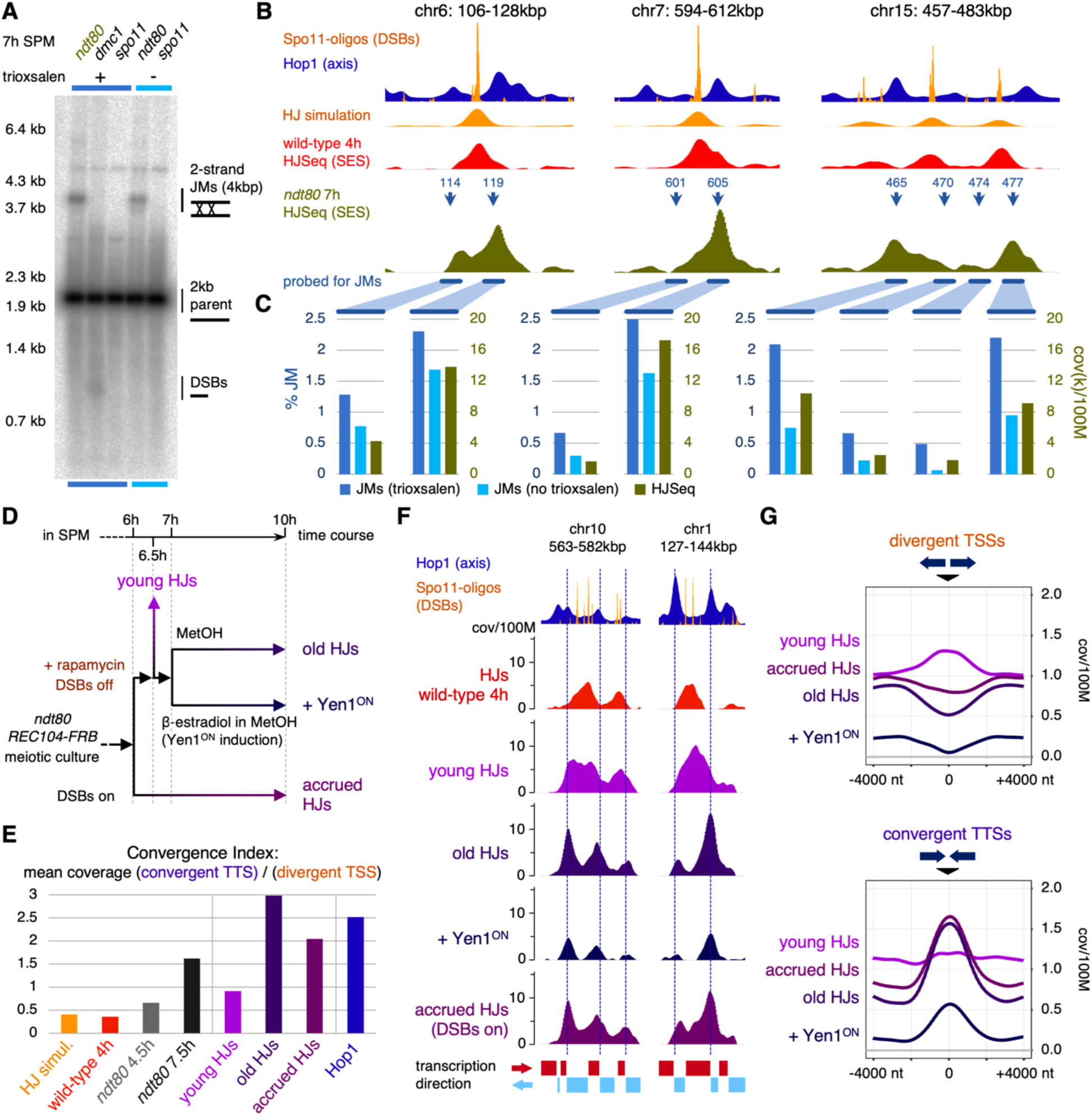
Holliday junctions branch migrate towards convergent transcription sites. (A) Example Southern blot joint molecule (JM) assay (non-covalently HJ preserving magnesium gel, locus chr15 915kbp). Parental DNA 2kbp, 2-strand JM ∼4kbp. Trioxsalen treated samples: *ndt80* 7h, *dmc1* 7h, *spo11 ndt80* 7h, used for HJSeq (SES). Uncrosslinked samples from an independent time course experiment: *ndt80* 7h and *spo11 ndt80* 7h. (B) Three regions with pronounced HJSeq patterns, mapped for JMs against HJSeq signals, *ndt80* 7h. Loci coordinates (in kbp) indicated (blue arrows). Tracks: Hop1 ChIP-Seq (blue, Shodhan et a. 2021) with Spo11-oligos (orange, Murakami et al. 2020), wild-type 4h HJ simulation (orange), wild-type 4h HJSeq (SES) (red), *ndt80* 7h HJSeq (SES) (olive, input median fragment size by weight 1.0kb). All Southern blot parental species are 1.9-2.2kb. (C) Quantification of *ndt80* 7h JMs (dark blue: trioxsalen treated, light blue: no trioxsalen) against HJSeq (SES) (olive) for loci in (B). JMs (2-strand) quantified against whole lane signal. Coverage: nucleotide coverage per 100M, in thousands. (D) Experimental setup for Spo11-DSB inhibition and Yen1^ON^ resolvase induction in *ndt80 REC104-FRB YEN1-IN* for calibrated HJSeq Query samples. (*spo11 ndt80 REC104-FRB YEN1-IN* experimental setup for Reference samples used in subtraction is outlined in Figure S4C). (E) Convergence Indices for the Spo11-DSB shutoff and Yen1^ON^ HJ resolvase induction experiment (*ndt80 REC104-FRB YEN1-ON*), ratio of mean coverage for convergent TTSs over mean coverage for divergent TSSs, genome-wide. (F) Example regions from *ndt80* Spo11-DSB shutoff Yen1-ON resolvase induction experiment. Calibrated HJSeq profiles, all samples of the DSB shutoff and Yen1^ON^ induction experiment were processed in parallel. Dashed lines indicate convergent TTSs. Red and blue rectangles: Transcription direction as per underlying ORF and co-oriented ORF runs (red: transcribed right-wards; blue: transcribed left-wards). (G) Genome-wide pile-up mean coverage analysis from Spo11 DSB shutoff and Yen1^ON^ HJ resolvase induction experiment for divergent TSSs (n=1417) and convergent TTSs (n=1413). Coverage is nucleotide coverage per 100M.

We also asked if the HJSeq signal that accumulates at convergent TTSs reflects the presence of branched DNA structures, using β-estradiol-induced expression of the engineered structure-selective endonuclease Yen1^ON^ ^7^, which cleaves a broad range of branched DNA structures including HJs ^42^. Expression of Yen1^ON^ after DSB depletion uniformly reduced HJSeq signals genome-wide by ∼73% over three hours (Figure 3F, 3G, S4G, and S4H), consistent with previous Southern blot-based measures of JM resolution upon DSB inhibition and Yen1^ON^ induction ^7^. Together, these data support a model in which HJs undergo directional branch migration during pachytene, progressively relocating the junctions towards convergent transcription sites.

### Directional HJ branch migration is enabled by the ZMM pathway

Crossover designated dHJs require the concerted activity of both the ZMM pathway and the Bloom’s helicase complex (in yeast STR: Sgs1-Top3-Rmi1). Moreover, STR targets nascent recombination intermediates and dHJs for disassembly to yield non-crossovers, unless protected by ZMM proteins (Figure 1A). Cells lacking STR therefore convert nearly all meiotic DSBs into undesignated dHJs that resolve equally into crossovers or non-crossovers ^7,8,43-51^. The branch migration necessary for intermediate dissolution is driven by the Bloom’s helicase Sgs1, while STR’s type-I topoisomerase Top3 relieves migration associated supercoiling ^37,49,52,53^. Thus, STR and ZMM proteins could influence bulk HJ redistribution during pachytene either by facilitating ATP-driven directed junction migration, or by restricting migration away from DSB sites via dissolution activity.

To examine the regulatory relationship between pachytene HJ branch migration, ZMM, and Bloom’s helicase activity, we first performed calibrated HJSeq in an *ndt80* strain that also carried the meiosis-specific Bloom’s helicase mutant *sgs1-mn* ^44^. At brief pachytene duration (4.5h), *sgs1-mn ndt80* showed the expected genome-wide increase in HJ levels (1.8-fold) relative to *SGS1 ndt80*. After another 3 hours in pachytene (7.5h), HJs accumulated further, to levels 3.7-fold higher than in *SGS1 ndt80*, which remained nearly unchanged (Figure 4C). These data indicate that STR is required to restrain excessive HJ-formation competence in the extended pachytene of *ndt80*.

**Figure 4.**
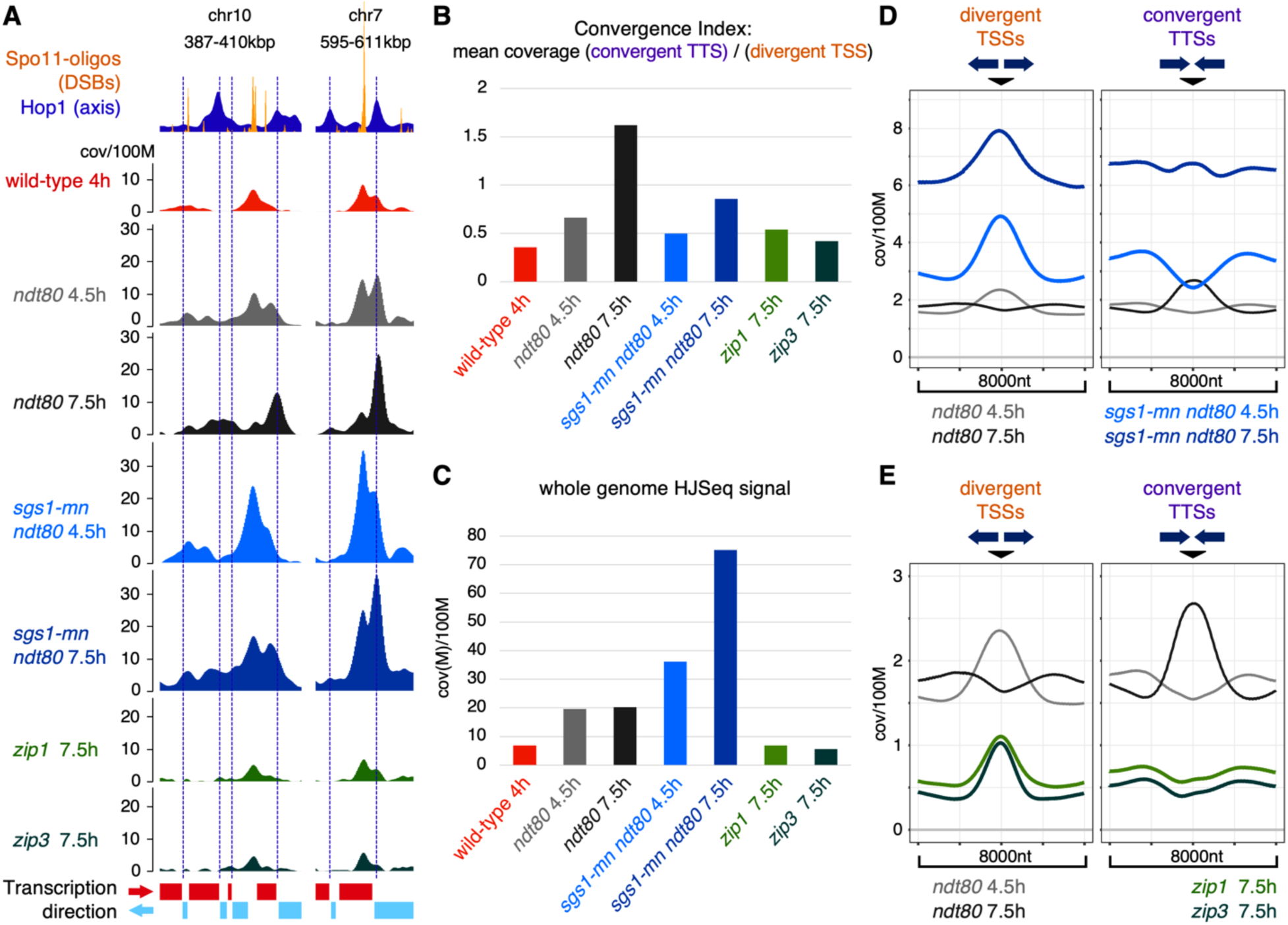
Directional long-range HJ migration is driven independently of Blooms helicase Sgs1, but depends on ZMM crossover control. (A) Two example regions of calibrated HJSeq from *sgs1-mn ndt80* (4.5h and 7.5h) and *zmm* mutants *zip1* and *zip3* (7.5h). Prophase I cells: *zip1* 7.5h: 77% (n=200), *zip3* 7.5h: 100% (n=200), by whole cell spindle immunostaining. Vertical dashed lines indicate convergent TTSs. Red and blue rectangles: transcription direction as per underlying ORF and co-oriented ORF runs (red: transcribed right-wards; blue: transcribed left-wards). Coverage: nucleotide coverage per 100M. (B) Convergence Indices for samples shown in (A), genome-wide mean coverage ratios of convergent TTSs signal over divergent TSSs signal. (C) Quantification of HJSeq (cal) signal genome-wide for experiments shown in (A, B). Coverage: nucleotide coverage per 100M, in millions. (D) Genome-wide pile-up mean coverage analysis of *sgs1-mn ndt80* (4.5h and 7.5h) against *ndt80* (4.5h vs 7.5h) for divergent TSSs (n=1419) and convergent TTSs (n=1414). Coverage is nucleotide coverage per 100M. (E) As in (D) for *zmm* mutants *zip1* and *zip3* (7.5h). Divergent TSSs n=1419, convergent TTSs n=1412.

Most importantly, HJ distribution patterns in *sgs1-mn ndt80*, while heterogenous, retained clear indications of migration during prolonged pachytene (Figure 4A, and S5). Genome-wide Convergence Index values, a measure of long-range HJ-relocation, increased substantially between 4.5 and 7.5h, albeit at 70% of the rate in *SGS1 ndt80* (Figure 4B). Pile-up analyses for *sgs1-mn ndt80* resolve these observations: under extended pachytene and increasing HJ levels, convergent TTSs gain substantially while HJSeq signals remain dominant at divergent TSSs (Figure 4D), consistent with ongoing formation and replenishment of HJs at divergent TSSs as “older” HJs migrated towards convergent TTSs.

We next examined the *zmm* mutants *zip1* and *zip3*. Zip1 is an essential component of the SC; the E3 ligase Zip3 mediates the initiation of synapsis between homolog axes at crossover designated intermediates ^54-57^. *zmm* mutants disrupt homolog synapsis, causing sustained DSB formation and a prolonged prophase I, but still form substantial quantities of dHJs and recombination products ^7,8,40,58,59^. At 7.5h after induction of meiosis, both mutants displayed markedly reduced genome-wide HJ levels relative to *ndt80* at 4.5h (*zip1*: 0.38×; *zip3*: 0.32×) (Figure 4C, and S2H), despite their prolonged prophase I (at 7.5h, *zip1*: ∼77%, *zip3*: ∼100% prophase I cells, n=200). Analysis of HJSeq signal patterns at 7.5h showed that HJs remain enriched at divergent TSSs in both *zip1* and *zip3*, with only minimal accumulation at convergent TTSs (Figure 4A, 4E, and S5). Accordingly, the Convergence Indices for both mutants remained below that of *ndt80* at 4.5h (Figure 4B). Thus, HJ migration is virtually absent from both of these *zmm* mutants. In summary, these findings indicate that both Zip1 and Zip3 (and by implication the entire ZMM pathway) provide competence for long-range HJ migration as part of regulated dHJ maturation. They also exclude the possibility that Bloom’s helicase Sgs1, which stabilizes the pachytene HJ landscape by restraining HJ accumulation, is the only activity driving directional branch migration during pachytene.

## DISCUSSION

Meiotic prophase I is a prolonged stage that integrates DNA-level recombination with chromosome reorganization to generate a selected set of crossovers, ensuring genetic reshuffling and accurate homolog segregation. Despite decades of work, the chromatin behavior of meiotic HJs – the recombination intermediates that give rise to crossovers – remained poorly resolved because there has been no approach to detect HJs directly and at high throughput. By developing HJSeq, a method that enables genome-wide mapping of HJs at high spatial resolution, we have unlocked a previously inaccessible dimension of homologous recombination intermediate dynamics. Applied across defined pachytene windows, this approach revealed extensive spatial reorganization of HJs during meiotic prophase I, establishing meiotic HJs as highly dynamic structures within chromatin.

At early stages, the genomic distribution of HJs closely reflects DSB-formation activity. This is evident from the strong genome-wide correlation between Spo11-oligo-based prediction (HJ simulation) and wild-type HJSeq profiles, indicating that the disposition for dHJ formation in wild type is largely determined by DSB hotspot strength. As pachytene duration increases, however, this local correspondence progressively breaks down, and we observe extensive directional redistribution of HJs along chromosomes. Remarkably, this redistribution follows the orientation of underlying transcriptional units, and HJs accumulate near sites of convergent transcription. Thus, dHJ intermediates are not fixed to their sites of formation, but can undergo pervasive and large-scale repositioning by branch migration.

Branch migration, the traversal of exchange points between recombination partners along the DNA, is integral to homologous recombination and dHJ formation ^1,20,60,61^. While theoretical considerations and in vitro experiments suggest that topological constraints should limit long-range movement of such structures along chromatinized DNA ^26,37,39^, our findings reveal that meiotic dHJs nevertheless undergo extensive directional movement during pachytene, implying mechanisms that relieve associated supercoiling and negotiate passage through nucleosomes. This long-range redistribution does not require the Bloom’s helicase Sgs1, a factor previously implicated in branch migration and disassembly of recombination intermediates ^7,8,20,37,43,48,50,51,61^. Instead, HJ relocation depends on the regulated maturation of recombination intermediates within the ZMM pathway, suggesting that association of dHJs with crossover-designation complexes permits long-range junction movement while preserving crossover fate.

The uniform alignment of HJ redistribution with the transcriptional landscape raises the possibility that transcription-associated chromatin dynamics contribute to this process. Translocation of RNA polymerase II generates torsional stress and is coupled to nucleosome remodeling. This could provide a physical driving force capable of promoting branch migration of recombination intermediates. Activities that clear topological constraints and relieve torsional stress associated with dHJ movement are likely to be required. Thus, topoisomerases, including the Top3-Rmi1 component of the Bloom’s complex, as well as chromatin remodelers such as the conserved nucleosome remodeler Chd1, represent attractive candidates for facilitating junction movement within chromatin. Top3 has recently been implicated in maintaining dHJ architecture ^62^ and has been suggested to track with recombination intermediates during pachytene ^63^; Chd1 has been linked to regulated meiotic crossover formation ^64^.

The core implication of these findings is that HJs are not merely formed and resolved, but their distribution is remodeled as pachytene progresses. This migration is pervasive and part of the regulated maturation of dHJs in the ZMM pathway, suggesting that the capacity for extensive branch migration is intrinsic to regulated dHJs and integral to normal crossover formation and positioning. Notably, branch migration has been considered a mechanism in models of regulated dHJ stabilization, as well as crossover resolution ^20,65^. Long-range branch migration, highlighting meiotic HJs as responsive chromatin participants, likely preserves crossover precursors in the face of active processes on the meiotic chromatin like transcription, until an adequate set of intermediates is secured. Additionally, regulated extension of pachytene and associated branch migration could allow for long-range homology sampling by heteroduplex formation to limit crossovers between non-allelic regions, since mismatch recognition is associated with dHJ rejection^20,66^.

Collectively, our findings reframe pachytene as an active period of intermediate remodeling and provide a mechanistic basis for how meiotic cells sustain long-lived recombination intermediates in a dynamic chromatin environment while preserving robust crossover outcomes. Beyond these biological implications, HJSeq establishes a broadly applicable approach for direct genome-scale analyses of HJ dynamics across biological systems, enabling comparative and mechanistic interrogation of genome stability in contexts where HJs are central repair intermediates, such as meiosis, somatic DNA repair, and replication stress.

## ACKNOWLEDGEMENTS

We are grateful to Yawen Bai and Bing-Rui Zhou for help with HPLC purifying biotinylated peptides, Franz Klein and Ed Louis for strains, Doris Chen for providing bioinformatic advice, Akira Shinohara for antibodies, Franz Klein, Alex Kelly, Anca Segall and members of the Lichten and Matos labs for helpful discussions, and Stewart Durell for Autodock Vina guidance. We acknowledge the Galaxy project, used for remapping Hop1 ChIP-Seq data. We thank the Center for Cancer Research Genomics Core at NCI and the VBCF NGS facility for high throughput sequencing. Peptide docking simulations used resources at the NIH HPC Biowulf cluster (https://hpc.nih.gov). This work was supported by the Intramural Research Program of the NIH through the Center for Cancer Research at the National Cancer Institute, H2020-EU Marie Skłodowska-Curie grant 847548, Austrian Science Foundation FWF SFB Meiosis - 8807-B, and the European Research Council ERC grant 101002629. The contributions of the NIH authors are considered works of the United States Government. The findings and conclusions presented in this paper are those of the authors and do not necessarily reflect the views of the NIH or the U.S. Department of Health and Human Services.

## AUTHOR CONTRIBUTIONS

MX conceived the project, developed HJSeq, developed and performed the bioinformatic pipelines and analysis, and performed all experiments except for the ndt80 7hrs Southern blots performed by AS, and the DSB shutoff Yen1^ON^ induction YEPA time course and controls performed by NK. ML and JM supervised and mentored the project. MX, JM, ML wrote the manuscript.

## FIGURES AND FIGURE LEGENDS

**Figure S1.**
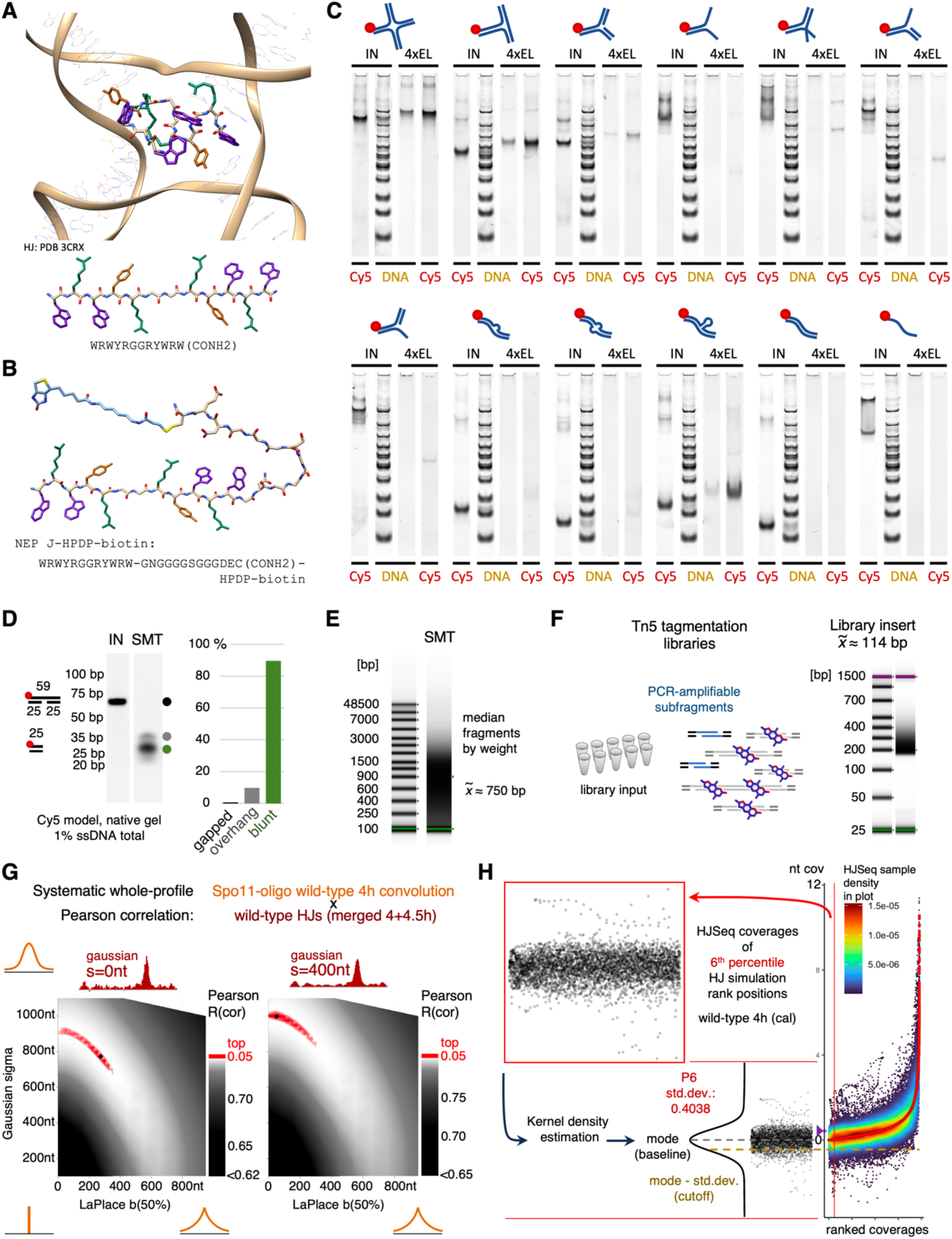
HJSeq method background, related to Figure 1. (A) Dodecameric HJ-binding peptide docked to an open-conformation HJ (AutoDock Vina generated, on PDB:3CRX) (top panel), and linear peptide (bottom panel). (B) NEP J-HPDP-biotin variant, derived from the dodecameric peptide, for HJSeq. C-terminal linker with final cysteine and conjugated HPDP-biotin (light blue). (C) HJ-binding peptide pulldown on Cy5-labelled model substrates, without prior DNA shearing and ssDNA digestion. Gel analysis for quantification shown in Figure 1B. Eluates loaded at 4-fold ratio relative to input. SSB binding to ssDNA reduces mobility. All species included in the quantification. Input and eluates were run on different gels. (D) SMT reaction for enzymatic shearing of genomic DNA and ssDNA/RNA degradation: 9nt gapped model substrate spike-in on native gel, used for calibrating Mung Bean Nuclease activity. ssDNA stretch of model at 1 percent of total DNA. Green: cut and blunted; gray: cut; black: full length gapped substrate parental species. (E) Genomic DNA fragment distribution after SMT reaction for wild-type 4h (cal). Median fragment size by weight ∼750bp. (F) Sequenceable subfragments are obtained by Tn5 tagmentation library preparation. DNA is ‘over’-tagmented for sub-fragments devoid of trioxsalen adducts. Example fragment distribution for Eluate wild-type 4h (cal) library shown. (G) HJ simulation obtained by convolution of wild-type 4h Spo11-oligo DSB coverage data (SRR7811345, Murakami et al. 2020) and systematic fitting by genome-wide Pearson correlation to wild-type (4+4.5h, cal) HJSeq profiles (left: raw HJSeq; right: denoised HJSeq data). To accommodate for both stochastic as well as exponential decay components in the observed HJSeq data, Spo11-oligo data was systematically convolved by Gaussian smoothing (sigma on y-axis) and LaPlace smoothing (b(50%) kernel spreading constant on x-axis). The respective Pearson correlation coefficient (R) is presented by color in the generated heat map. The black cross indicates R(max): 0.74506 left, 0.80471 right. Spo11-oligo profiles smoothed by LaPlace b50=275nt and Gaussian s=775nt describe raw wild-type HJ profiles best. For standard denoised (Gaussian s=400nt smoothed) HJSeq profiles, the profile from Gaussian smoothed s=1000nt Spo11-oligo data is used as ‘HJ simulation’. (H) Schematic of zero baseline adjustment and noise estimation on HJSeq profile coverages, after Query-Reference subtraction, on the coldest 6% meiotic recombination regions in the genome as per wild-type HJ simulation. Example for wild-type 4h (cal) HJSeq data (denoised: Gaussian s=400nt, 150nt bins, region exclusions applied) is shown. Single nucleotide coverages are sampled from all HJ simulation (normalized to total HJSeq coverage) and HJSeq 150nt bins genome-wide. Both coverage datasets are ordered (i.e. ranked) by the HJ simulation (Spo11 DSB-based) signal, while preserving positional correspondence between datasets. Consequently, ranked HJ simulation coverages follow a monotonic pseudo-exponential progression (reflecting the Spo11-DSB hotspot strength distribution over the genome), while HJSeq coverages at these respective positions disperse around the HJ simulation coverages. This correspondence between HJ simulation and HJSeq coverages is captured in ranked HJSeq coverage heat dot plots (right). Red with black outline: HJ simulation coverages. Heat dot plot: HJSeq coverages following a linear color scale that indicates their local density in the plot. X-axis indicates the ordered nucleotide coverage sample progression (n > 8000 at 150nt bins), y-axis indicates the nucleotide coverage value (as coverage per 100M) adjusted to the estimated true baseline. Purple tick: original zero baseline after subtraction of scaled *spo11* Reference from scaled Query. Red vertical line: Border of the 6^th^ percentile lowest coverage regions in the HJ simulation (P6), used for baseline and noise estimation. For estimating the true zero baseline and the signal variation in the HJSeq data, the HJSeq signals in the coldest 6% meiotic recombination regions (P6) is analyzed (top left inset, red outline). Bottom, left: The adjusted HJSeq zero baseline is estimated by finding the mode in the P6 HJSeq coverages by kernel density estimation (black, Sheather Jones bandwidth*5) and identifying its maximum (‘P6 mode’, gray dashed line). The standard deviation is calculated (‘P6 std.dev.’), and the lower cutoff for pile-up heatmaps is determined as ‘P6 mode - P6 std.dev.’ (gold dashed line).

**Figure S2.**
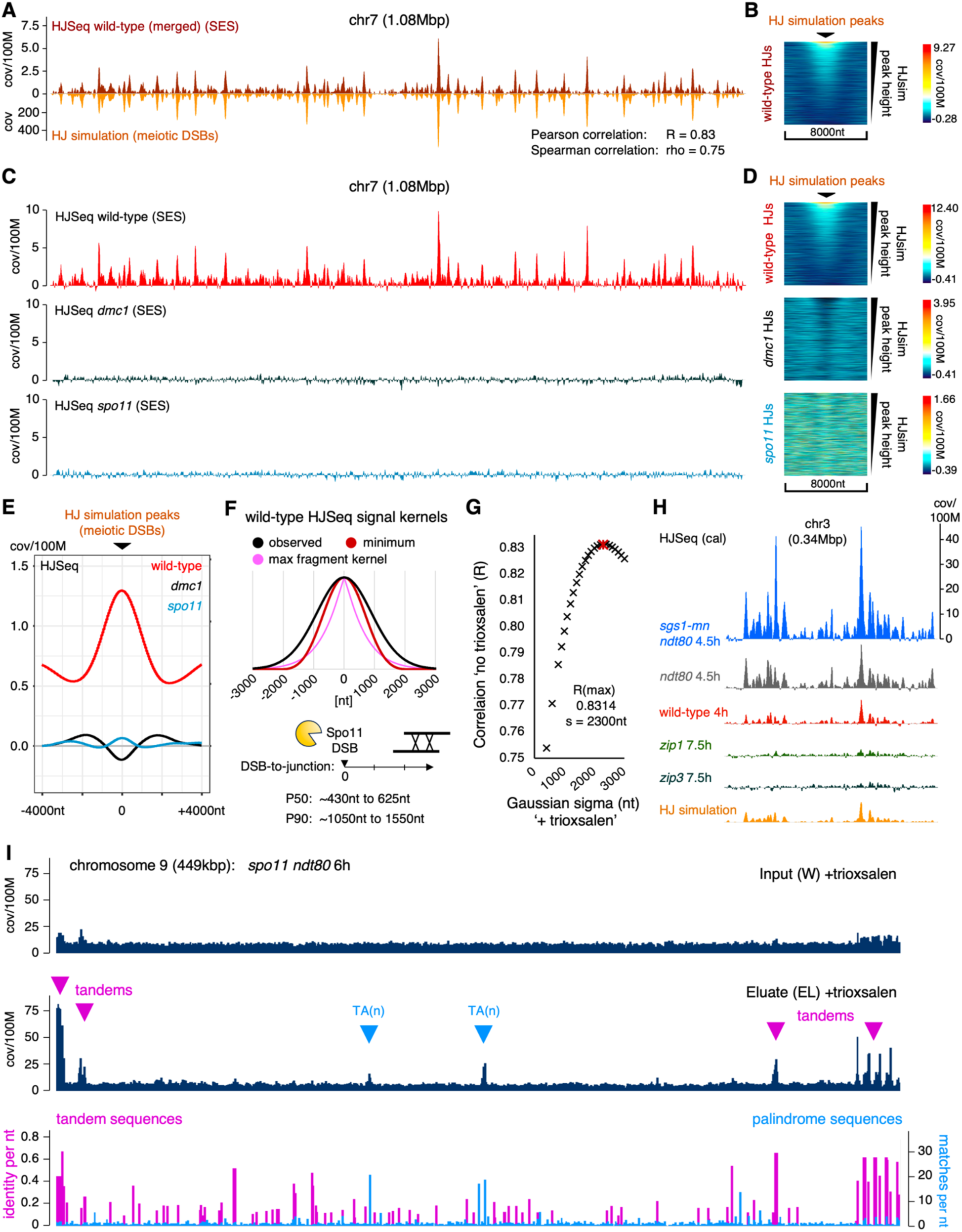
Meiotic HJSeq signals report HJ intermediates, related to Figure 1. (A-E) Biological replicate HJSeq set (not calibrated, SES) for wild type, *dmc1*, and *spo11*. (A) Chromosome 7 (1.08Mbp), wild-type 4+4.5+5h HJSeq (averaged, SES) (top, dark red, 150nt binning, denoised) against the HJ simulation (bottom, orange, as in Figure 1E) with y axis scale normalized to total wild-type HJSeq signal. Single nucleotide coverage is reported. (B) Genome-wide pile-up heatmap for wild-type 4+4.5+5h HJSeq (averaged, SES) coverages at HJ-simulation peak positions (n=2118). Linear color-to-coverage relation as indicated, scaled to coverage range. Lower cutoff applied. Coverage: single nucleotide coverage per 100M. (C) Light red: wild-type 4h (SES). Black: *dmc1* 5h (SES), turquoise: *spo11* 4.5h (SES). Whole chromosome 7 HJSeq profiles shown. (D) Genome-wide pile-up heatmaps for HJSeq data of (C) for HJ-simulation peak positions. Linear color-to-coverage relation as indicated, scaled to coverage range. Lower cutoff applied. Coverage: single nucleotide coverage per 100M. (E) Genome-wide mean coverage pile-up analysis relative to HJ-simulation peak positions on data shown in (D). Coverage: single nucleotide coverage per 100M. (F) Average wild-type HJSeq signal footprint from DSB position. Black: Best convolution fit of raw Spo11-oligo (DSB) profile to observed wild-type HJSeq profile (4+4.5h, calibrated, not denoised), using Gaussian and Laplace components (see Figure S1G). Pink: Maximum fragment distribution kernel (weighted by sample DNA mass distribution, scoring 250-3000bp, 87% of input sample fragment distribution). Red: Minimum average HJ kernel approximation by deconvolution of observed with fragment distribution kernel. Kernels are displayed normalized to summit height. Distribution percentiles (P50: 50%, P90: 90%) of the approximated minimum and the observed HJ kernel are indicated as DSB to (outer) junction span. (G) HJSeq signals broaden in absence of trioxsalen interstrand crosslinks. Pearson correlation fitting of a regular (+trioxsalen) HJSeq profile, Gaussian smoothed at indicated bandwidths, to the corresponding ‘no trioxsalen’ HJSeq profile. Samples were obtained and treated in parallel throughout (*ndt80* 6h, calibrated). Pearson R(max) of series indicated in red (R=0.8314, s=2300nt). (H) Testing genetic predictions for meiotic dHJs with calibrated HJSeq. In scale HJSeq chromosome 3 profiles for indicated mutants and wild type. HJ-simulation profile (orange) provided as reference for recombination hotspots. *zmm* mutants are prophase I arrested (*zip1* 7.5h: 77% (n=200) of cells in prophase I, *zip3* 7.5h: 100% (n=200); by whole cell spindle immunostaining). (I) Tandem direct repeat sequences with substantial self-pairing ability and TA-run (n≥10) palindromic sequences form meiotic recombination independent signals. Input and HJ-pulldown eluate profiles (i.e. unsubtracted) from *spo11 ndt80* 6h, whole chromosome 9 rich in tandem and TA(n) regions. Triangles indicate pronounced eluate signal independent of recombination (magenta: tandems; blue: TA(n) palindromes). Bottom profile: tandem repeats (magenta) and palindromes (blue) scored with EMBOSS etandem (≥4 16-170nt repeats, consensus match-mismatch score per nt in region, maximum is 1) and EMBOSS palindrome finder (12-100nt palindromes, max gap 170nt, max mismatch 2, allowing overlaps, repeat coverage per nucleotide) on the SK1 ASMv1 reference sequence. Profile binning of 25nt was applied on the resulting scores. Y-axis: average score per bin.

**Figure S3.**
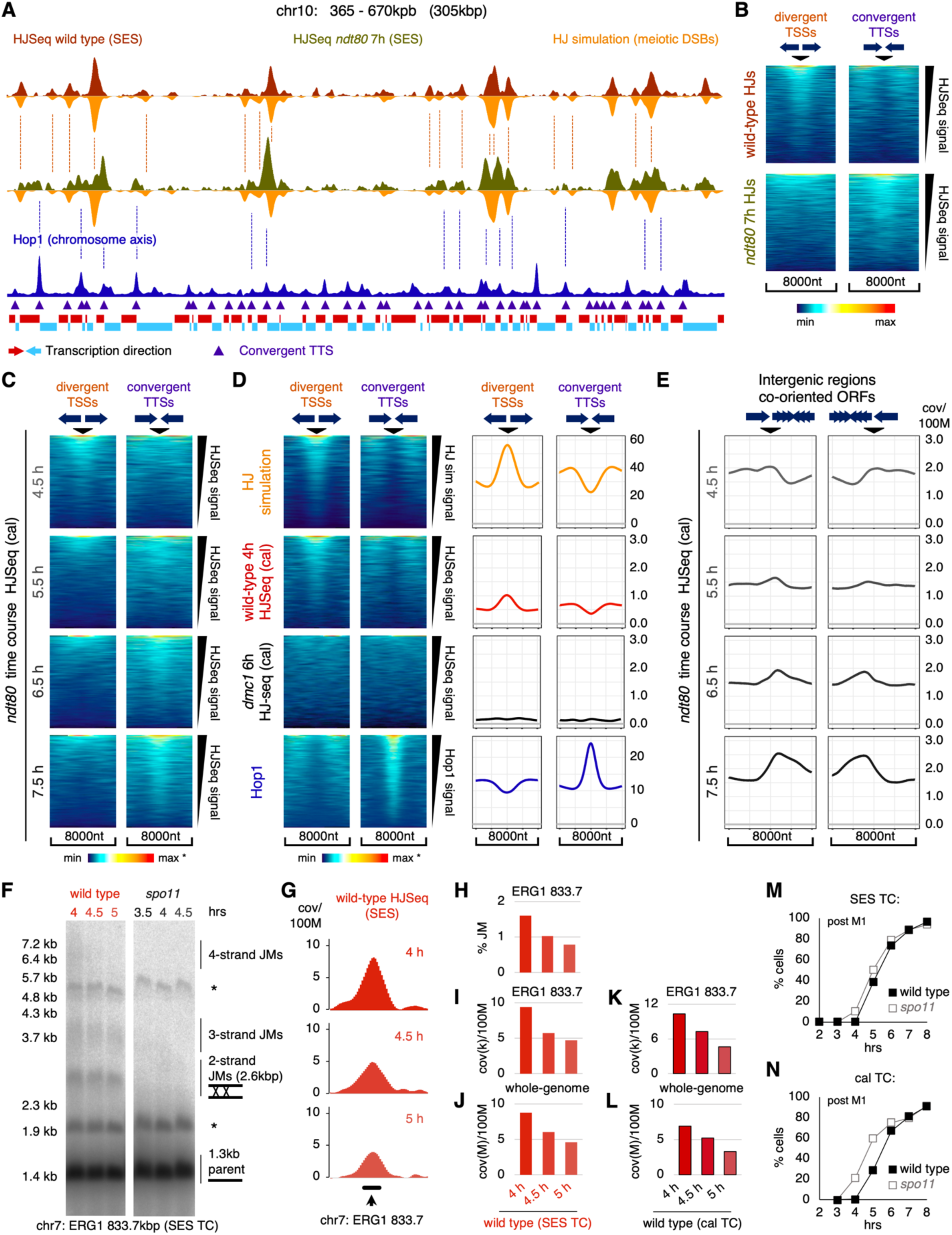
Related to Figures 2 and 3. (A) Biological replicates to Figure 2A. Red: HJSeq wild-type 4+4.5+5h (SES, averaged). Olive: Pachytene arrested *ndt80* 7h (SES, input median fragment size by weight 1.0kb). Orange: HJ simulation (from Spo11-oligos wild-type 4h, Murakami et al. 2020). Blue: Hop1 ChIP-Seq (Shodhan et al. 2021), indicating chromosome axis sites. Violet triangles: Convergent TTSs. Red and blue rectangles: Transcription direction as per ORF and co-oriented ORF runs (red: transcribed right-wards; blue: transcribed left-wards). (B) Genome-wide pile-up heatmap analysis of HJSeq data of (A) for divergent TSS (n=1419) and convergent TTS positions (n=1411) (respective center points) over 8kbp regions. Sorted by HJSeq signal per region. Heatmap colors scaled to coverage range in heatmap. Coverage follows heat scale linearly. Lower cutoff applied to coverages for heatmaps. (C) Genome-wide pile-up heatmaps for HJSeq (cal) coverages from *ndt80* mutant time course series at divergent TSS (n=1419) and convergent TTS (n=1414) positions (of Figure 2D). Coverage follows linearly the color scale. Heatmap color range is scaled to minimum and maximum coverage. Coverage ranges (min,max; nt cov/100M): 4.5h Divergent -0.61, 24.71; 4.5h Convergent -0.61, 23.32; 5.5h D -0.71, 27.10; 5.5h C -0.71, 17.84; 6.5h D -0.97, 31.25; 6.5h C -0.97, 20.35; 7.5 D -1.37, 37.28; 7.5 C -1.37, 24.96. Lower coverage cutoff was applied for heatmaps. (D, left) Genome-wide pile-up heatmaps at divergent TSS (n=1419) and convergent TTS (n=1414) positions for HJ simulation (meiotic DSBs, wild-type 4h), HJSeq wild-type 4h (cal), HJSeq *dmc1* 6h (cal), Hop1 ChIP-Seq (wild-type 4h, Shodhan et al. 2021). Coverage follows linearly the color scale. Heatmap color range is scaled to minimum and maximum coverage of regions. Coverage ranges (min,max; HJ simulation and Hop1 is nt cov; HJSeq is nt cov/100M): HJ simulation D 0.0, 581.7; HJ simulation C 0.0, 580.8; HJSeq WT 4h D -0.40, 11.39; HJSeq WT 4h C -0.40, 11.39; HJSeq *dmc1* 6h D -0.54, 9.70; HJSeq *dmc1* 6h C -0.54, 9.70; *Hop1* WT 4h D 0.3, 114.5; *Hop1* WT 4h C 0.6, 128.3. Lower coverage cutoff was applied for HJSeq heatmaps. (D, right) Genome-wide pile-up mean coverages at divergent TSS (n=1419) and convergent TTS (n=1414) positions for HJ simulation (meiotic DSBs, wild-type 4h), HJSeq wild-type 4h (cal), HJSeq *dmc1* 6h (cal), Hop1 ChIP-Seq (wild-type 4h, Shodhan et al. 2021), as in (D). Coverage HJSeq: nucleotide coverage per 100M. Coverage HJ simulation and Hop1: absolute nucleotide coverage. (E) *ndt80* time course (4.5-7.5h) genome-wide pile-up mean coverage analysis for relocation of HJSeq (cal) signal at regions of co-oriented ORFs. Regions for FWD-(Center)-FWD-REV (n=648) and FWD-REV-(Center)-REV (n=634) ORF configurations are shown. Convergent positions distribute due to variable ORF lengths. (F) Southern blot joint molecule assay on trioxsalen treated genomic DNA from wild type and *spo11* (HJSeq, SES, Query and Reference samples). Non-covalently HJ-preserving magnesium gel. Probed for a 1.3kb fragment (EcoRV-XhoI) at the strong endogenous recombination hotspot ERG1 (chr7 833.0 - 834.3 kbp). Time points and fragment species as indicated. Asterisks indicate trioxsalen partial restriction digestion fragments (approximately 1.8kb and 5.0kb). (G) ERG1 hotspot region HJSeq (SES) profiles, wild-type 4, 4.5 and 5h using *spo11* 3.5, 4 and 4.5h as subtraction Reference, respectively. From genomic DNA as in (F). Region probed in (F) indicated by black bar. (H) Joint molecule quantification of data shown in (F), scoring 2-strand, 3-strand, and 4-strand (all) JMs relative to the whole lane signal, for indicated time points. (I, J) HJSeq (SES) quantification for wild-type 4, 4.5 and 5h ERG1 EcoRV-XhoI fragment region (F, H), and genome-wide. Coverage is nucleotide coverage per 100M, in thousands or millions respectively. (K, L) HJSeq (cal) quantification for wild-type 4, 4.5 and 5h ERG1 EcoRV-XhoI fragment region, and genome-wide. Coverage is nucleotide coverage per 100M, in thousands or millions respectively. (M, N) Meiotic progression for the HJSeq ‘SES’ time course (M) and ‘cal’ time course (N) cultures (wild type and *spo11*). Meiotic nuclear divisions on ethanol fixed, DAPI stained cells. Percentage of cells past metaphase 1 is shown (n=200 scored cells per time point).

**Figure S4.**
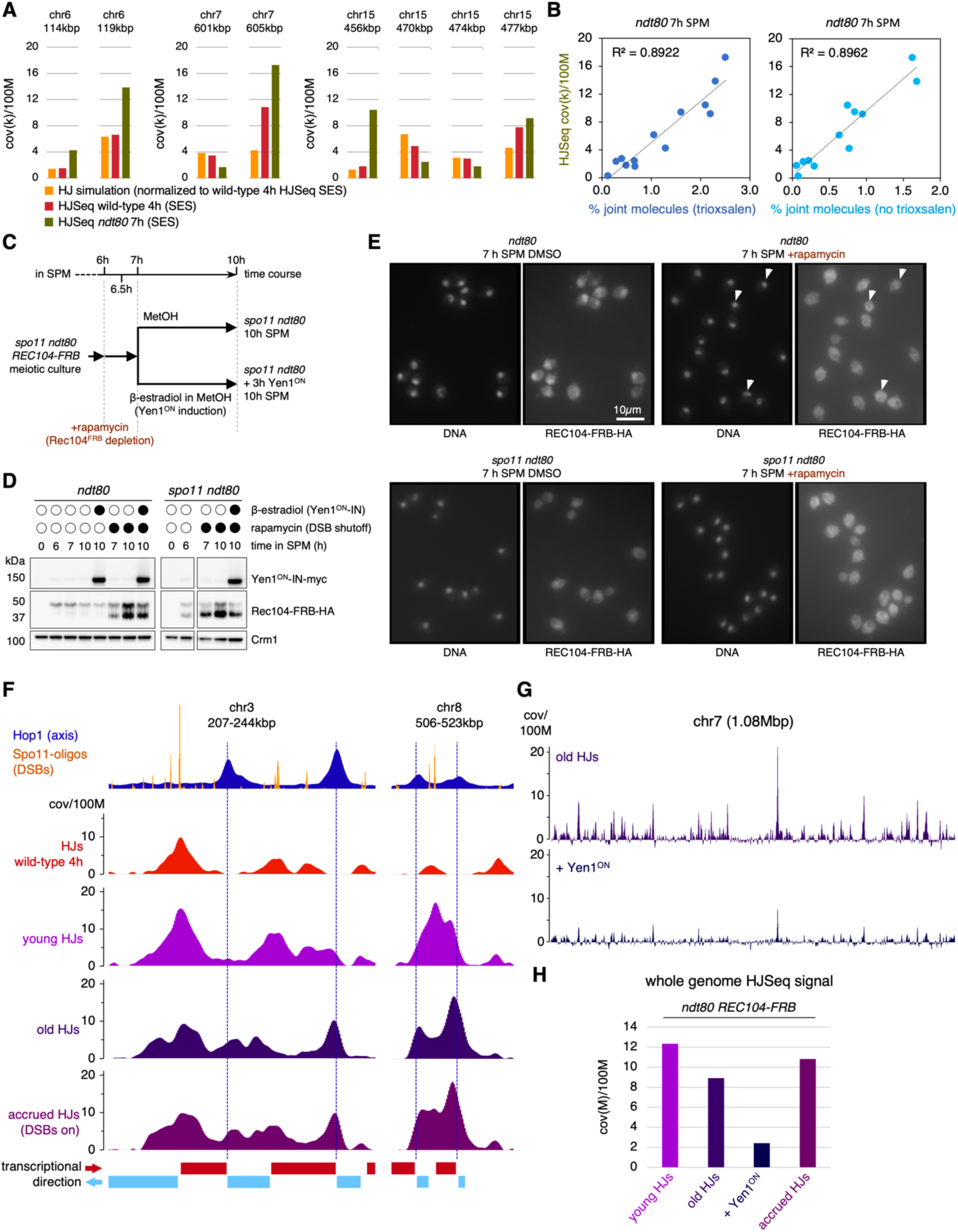
Related to Figure 3. (A) Related to Figure 3C. Profile quantifications at loci shown in Figures 3B and 3C: HJ simulation, wild-type 4h HJSeq (SES), *ndt80* 7h HJSeq (SES, input median fragment size by weight 1.0kb). HJ simulation data was normalized to total genome coverage of wild-type 4h HJSeq (SES). Nucleotide coverage per 100M, in thousands. (B) Bivariate plots comparing Southern blot joint molecule quantification against *ndt80* 7h HJSeq (SES) signal. Left: JMs from trioxsalen treated genomic DNA used for HJSeq. Right: JMs from independent time course experiment (not trioxsalen treated). Dotted lines: Linear regression. Loci as in (B, C), as well as chr3:325kb, chr9:106kb, chr9:150kb (trioxsalen treated only), chr15:846, chr15:915. Quantification: %JMs relative to lane signal, and nucleotide coverage per 100M, in thousands. All Southern blot parental species are 1.9-2.2kb. (C) Experimental setup for obtaining *spo11 ndt80* Reference HJSeq samples in Spo11-DSB inhibition Yen1-ON induction experiment (used in subtraction for differential HJSeq profiles with Queries obtained as in Figure 3D). (D) Western blot for Rec104-FRB-HA and inducible Yen1^ON^-myc. Loading control Crm1. Controls for the Spo11-DSB inhibition and Yen1-ON induction experiment. Lower band for Rec104-FRB-HA putative accumulation of unphosphorylated Rec104 protein outside of the nuclear environment and the Spo11 break-machinery (Kee et al., 2004). (E) Whole cell in situ immunofluorescence control for cellular localization of Rec104-FRB-HA (all images same scale). Controls for the Spo11-DSB inhibition Yen1-ON induction experiment. Arrows indicate example nuclei in *spo11*. (F) Two additional example regions for the *ndt80 REC104-FRB* experiment, promiscuous for HJSeq signal accumulation when pachytene DSB formation was permitted. Dashed lines indicate convergent TTSs. Red and blue rectangles: Transcription direction as per underlying ORF and co-oriented ORF runs (red: transcribed right-wards; blue: transcribed left-wards). Coverage: nucleotide coverage per 100M. (G) Whole chromosome 7: ‘old HJs’ (top) and ‘+Yen1^ON^‘ (bottom) HJSeq profiles, indicate homogenous Yen1^ON^ sensitivity of HJSeq signals. Coverage: nucleotide coverage per 100M. (H) Quantification of HJSeq (cal) signal genome-wide for Spo11-DSB inhibition Yen1^ON^ induction experiment. Coverage: nucleotide coverage per 100M, in millions.

**Figure S5.**
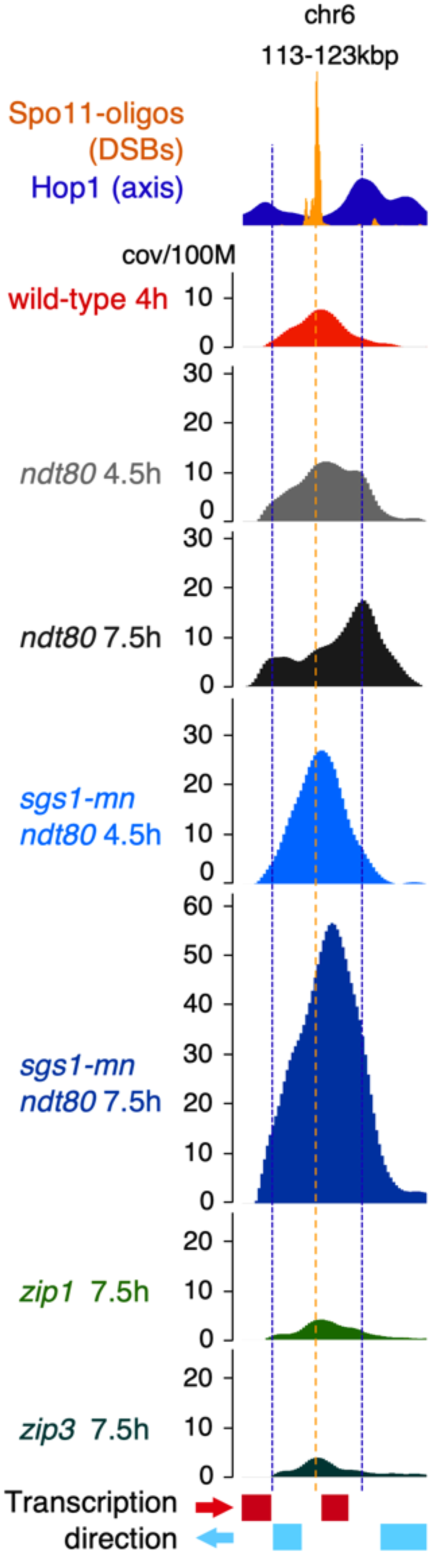
Example locus on chr6 (113-123kbp) with strong HJSeq signal accumulation in *sgs1-mn ndt80*, related to Figure 4. Calibrated HJSeq from *sgs1-mn ndt80* (4.5h and 7.5h) and *zmm* mutants *zip1* and *zip3* (7.5h). Blue vertical dashed lines indicate convergent TTSs. Orange vertical dashed line marks predominant Spo11 DSB position. Red and blue rectangles: transcription direction as per underlying ORF and co-oriented ORF runs (red: transcribed right-wards; blue: transcribed left-wards). Coverage: nucleotide coverage per 100M.

## SUPPLEMENTAL INFORMATION

### METHODS

#### Yeast strains

All strains used in this study are derivatives of SK1. Detailed genotypes are provided in Table S1. Strains were constructed by crossing or LiAc transformation, using standard procedures. All alleles have been described previously. For *sgs1-mn*, the endogenous promoter is replaced by a CLB2 promoter fragment ^44^. The *REC104-FRB-HA* anchor away and estrogen inducible *YEN1^ON^-IN-myc* system has been described ^7,41^. *ndt80*, *dmc1*, *zip1* and *zip3* are ORF deletions, *spo11* is either the catalytically dead allele (Y135F), or an ORF deletion (Rec104^FRB^ anchor away, Yen1^ON^-IN-myc system).

#### Meiotic time courses - SPS method

All experiments including *S.mikatae* meiotic spike-in used this method, except for *REC104-FRB-HA* anchor away and inducible *YEN1^ON^-IN-myc*. Diploid yeast strains were synchronized for meiosis as described previously ^67,68^. In brief, cells from a single colony were grown 24 hours at 30°C in liquid YPD (1% Difco Bacto yeast extract, 2% Difco Bacto peptone, 2% dextrose, pH 5.5). For synchronous meiosis in liquid culture, SPS pre-sporulation media (0.5% Difco Bacto yeast extract, 1% Difco Bacto peptone, 0.17% Difco Bacto yeast nitrogen base w/o AA, 1% potassium acetate, 0.5% ammonium sulphate, 0.05 M potassium-biphthalate, pH 5.5) was inoculated from the YPD culture at 1:1500-1:2500 and grown overnight at 30°C to an OD600 of 1.2-1.3. Cells were collected by centrifugation and washed once with prewarmed sporulation media (SPM: 1% potassium acetate, 0.001% PPG2000, and depending on auxotrophies with 4 mg/L Ura, 4 mg/L Trp, 4 mg/L His, 4 mg/L Arg, 6 mg/L Leu). The cells were then resuspended in prewarmed SPM to approximately 3-4x10^7^ cells/mL, marking time point 0h SPM, and incubated in baffled 2.8L Fernbach flasks at ≥300rpm for high aeration at 30°C.

#### Meiotic time courses - YEPA method

Utilized for the Rec104^FRB^ anchor-away and Yen1^ON^ induction experiment; as described previously ^7,62^. In brief, a single colony preselected on YPG plate (1% Difco Bacto yeast extract, 2% Difco Bacto peptone, 2% glycerol, 2% Difco Bacto agar) at 30°C for 48h, is patched on YPD plate (1% Difco Bacto yeast extract, 2% Difco Bacto peptone, 2% dextrose, 2% Difco Bacto agar) grown for 24h 30°C, then patched to a lawn on YPD plates and expanded for another 24h 30°C. YEPA (aka YPA) pre-sporulation liquid media (1% Difco Bacto yeast extract, 2% Difco Bacto peptone, 2% potassium acetate) was inoculated with the obtained cells to OD600 0.3 and grown 14h into a G1 arrest at 25°C. Cells were washed with prewarmed sporulation media SPM (2% potassium acetate), and resuspended in pre-warmed SPM to approximately 5-7x10^7^ cells/mL. Pre-sporulation and sporulation cultures were performed in an aerated 10L fermenter system. Sufficient Antifoam A concentrate (Sigma A6582) was added during pre-sporulation to prevent foam layer formation. Cultures were split into 5L Erlenmeyer flasks to a 30°C shaker for rapamycin (Sigma 553211, 4.6ug/uL in DMSO, to 1ug/mL in SPM) or DMSO mock addition at 6h SPM (Rec104^FRB^ anchor-away), and β-estradiol (Sigma E8875, 5mM in methanol, to 1uM in SPM) or methanol mock addition at 7h SPM (pGAL1-YEN1^ON^, via GAL4(848).ER ^69^).

#### DNA cytometry

Cellular DNA content was assayed from 0h SPM to 4h (SPS) or to 5-6h (YEPA) to verify progression through pre-meiotic S-phase. Cells from 1mL meiotic culture were fixed in 70% ice cold ethanol and stored at 4°C or ice until processed. Cells were washed in 1mL, then resuspended in 0.5mL 50mM Tris pH 7.5. RNA was digested with 200ug RNase A (Qiagen 19101) at 37°C for 2-4h. Cells were washed in 1mL FACS buffer (200mM Tris pH 7.5, 211mM NaCl, 78mM MgCl_2_), then resuspended in 1mL FACS buffer with 25ug/mL propidium iodide and briefly sonicated. 50ul of sample were transferred to 1mL 50mM Tris pH 7.5 for cytometry, performed on a FACSCalibur (BD Biosciences). Data were analyzed with FlowJo.

#### Cytology

Yeast cytology was performed as previously described ^67^. Meiotic nuclear divisions were followed on 70% ethanol fixed, then DAPI (0.2ug/mL in water) stained whole cells. For whole cell in situ immunostaining, formaldehyde was added on 1.5mL SPM culture to 3.2%, incubated overnight at 4°C. Cells were washed three times, and cell walls digested 15-40min in 150ul 1M Sorbitol, 2% potassium acetate with 70mM DTT and 125ug Zymolyase 20T (Nacalai) at 37°C. Cell suspension was mounted on poly-L-lysine coated multi-well slides, fixed 3min in methanol and 10sec acetone (ice cold). Cells were blocked 10min and immunostained 2-3h RT or overnight 4°C per primary and secondary antibody step in 0.5% BSA fraction V and 0.2% gelatin in 1xPBS. Spindle and Rec8 staining: rat, anti-tubulin 1:1000 (abcam 6160) and rabbit, anti-Rec8 serum 1:1000 (gift from Akira Shinohara), donkey, anti-rabbit-Cy3 1:800 (Jackson 711-165-152), rabbit, anti-rat-FITC 1:600 (Sigma F1763). Rec104-FRB-HA staining: mouse, anti-HA 16B12 1:200 (BioLegend 901501), goat, anti-mouse-Alexa 488 1:300 (Invitrogen A11029). Vectashield with DAPI (Vector Labs H-1200) or ProLong Diamond Antifade Mountant with DAPI (Invitrogen P36966) was used to stain DNA and stabilize the preparation. Whole cell IF images were taken on a Zeiss Axio Imager M2, 63× oil-immersion Plan-Apochromat objective (1.4 NA), CoolSNAP HQ2 camera, Zeiss ZEN blue 3.3. Chromosome surface spreads from *S.mikatae* were obtained from 1.5mL SPM culture collected at 4h15min (spike-in sample collection time). Cell walls were digested in 100ul 1M Sorbitol, 2% potassium acetate, 16mM DTT, 30ug Zymolyase 100T (Nacalai) at 30°C. Spheroplasting was followed by phase contrast microscopy, tested by addition of Sarkosyl to 0.5%. 6ul spheroplasts were mixed with 60ul 2.4% paraformaldehyde, 1.9% sucrose, 0.38% Lipsol on a slide, gently streaked, and allowed to dry at RT. Slides were washed in 1xPBS, and chromatin spreads were blocked 10min and immunostained 2-3h RT or overnight 4°C per primary and secondary antibody step in 0.5% BSA fraction V and 0.2% gelatin in 1xPBS. Zip1 staining: rabbit, anti-Zip1 1:480 (custom made, recombinant S.c. Zip1 1-227 his6), donkey, anti-rabbit-Alexa 488 1:500 (Invitrogen A21206). Vectashield with DAPI (Vector Labs H-1200) was used to stain DNA and stabilize the preparation. Imaging was performed on a Zeiss Axioskop, 100x oil-immersion Plan-Neofluar objective (1.3 NA) 2.5x Optovar, Photometrix Quantix camera and Molecular Devices MetaMorph.

#### Western blotting

Protein analysis was performed as previously described ^7,62^. In brief, cells (9mL SPM) were pelleted (3min, 860rcf, 4°C), resuspended in cold 10% TCA, and glass beads lysed via a FastPrep-24 5G (MP Biomedicals; two 40sec cycles at 6m/sec, 4°C). Precipitates were collected (10min, 21130rcf, 4°C), resuspended in 2x NuPAGE sample buffer with 200mM DTT, and acid neutralized adding 0.5x vol 1M Tris. After boiling (10min, 95°C), debris was cleared (10min, 21130rcf), and the supernatant was collected. For quantification, Bio-Rad protein assay was used. Samples were resolved on NuPAGE 3-8% Tris-Acetate gels (Invitrogen) and transferred to 0.45um PVDF membranes (Amersham Hybond). Immunostaining primary antibodies: rabbit, anti-Myc-HRP 1:10000 (Abcam ab1326), mouse, anti-HA 16B12 1:5000 (BioLegend 901501), rabbit, anti-Crm1 1:5000 (gift from Karsten Weis). Secondary antibodies: sheep, anti-mouse-HRP 1:10000 (Cytiva NA931), donkey, anti-rabbit-HRP 1:10000 (Cytiva NA934). Image acquisition via the Bio-Rad ChemiDoc MP Imaging System.

#### Physical JM analysis by Southern blotting

Genomic DNA was obtained using a CTAB extraction procedure as previously described ^70^ with modifications as per Detailed HJSeq Workflow section. Electrophoresis under HJ preserving conditions, alkaline Southern blotting to Amersham Hybond XL, and SSPE hybridization were performed as described ^70^. Imaging was performed with an Amersham Typhoon biomolecular imager. Assayed loci, restriction fragments and probes are provided in Table S3. JM species (2-strand JMs and mcJMs wild-type 4h; 2-strand JMs *ndt80* 7h) were quantified against whole lane signal using Fuji Image Gauge 4.22.

#### Peptides

Peptides (L-amino acids) were synthesized by New England Peptide to 1mg net by AA content to ≥95% purity, C-terminus amidated, and were dissolved 4-5min at RT in DMSO (≥99.9%, anhydrous, Sure/Seal, Sigma 276855-100ML) for 10mM master stocks, stored at -80°C. HJSeq “NEP J”: WRWYRGGRYWRWGNGGGGSGGGDEC-amide, carrying HPDP-biotin. Peptide structure illustration and visualization were performed with OpenChemistry Avogadro 2 and UCSF Chimera.

#### Model substrates and polyacrylamide DNA electrophoresis

A list of model substrates is provided in Table S2. The 5’ terminus of one oligo was fluorophore labelled with Cy5. Unlabeled oligos either PAGE or IE-HPLC purified, labelled oligos by PAGE or RP-HPLC, 20uM stocks in 0.3xTE. Annealing was performed at 2pmol/ul product in 1xAnneal Buffer (10mM Tris pH 8.0, 30mM NaCl). Native PAA gel electrophoresis was performed on 8%T, 3.3%C mini gels (1xTBE) at 110V for 50-70min RT. Gels were washed 15min in 1x running buffer before imaging fluorescently labeled substrates (Amersham Typhoon or Fuji FLA 5100). Afterwards, gels were stained with 1x SYBR Gold (in 1xTBE) for 15-25min, and washed twice for 5min (1xTBE), prior imaging gels again for total DNA.

#### Minimum average wild-type HJSeq distribution modelling

Observed wild-type HJSeq signals, i.e. a convolved signal in respect to punctuate HJs on pulled-down (sub)fragments, were described as a Gaussian and Laplace compound distribution of wild-type 4h Spo11-oligo peaks ^21^ (Figure S1G), based on best whole profile Pearson R between the raw HJ signal and a systematic series of smoothed Spo11-oligo profiles as the ‘observed wild-type HJ’ signal kernel. An approximation for the ‘maximum fragment contribution’ kernel was obtained using *spo11* 4hrs (cal) input TapeStation HS D1000 fragment mass distribution data 250-3000bp (87% of signal). Fragment sizes were extrapolated using the included MW size standard (9 size marker bands; max. deviation from MW standards: 11% at 3000bp, scored as 3319bp; less than

+/- 5% between 250-2500bp containing 85% of signal), and the TapeStation electrophoresis mass data converted to a linear single nt range representation of mass over the fragment size range (mass signal of single data points was distributed equally over each respective extrapolated range). For building a cumulative point signal kernel (limits +/-5000nt), uniform fragment availability for the pulldown with direct proportionality between DNA mass and probability for containing HJ DNA was assumed for simplicity, ignoring observed decreasing pulldown efficiency for larger fragments (therefore, and intentional overestimate of the fragment contribution to the observed signal). Deconvolution of the ‘observed’ average wild-type HJSeq peak distribution kernel with the approximate ‘maximum fragment contribution’ kernel (both +/-5000nt) was performed by FFT Fourier domain inverse filtering (Wiener) with empirical Tikhonov regularization in R: MIN <- OBS * ((Conj(FRAG) / (Mod(FRAG)^2)) + 0.000001), and the central +/-3000nt were kept. Validity of the kernel was empirically validated by convolution, closely reproducing the ‘observed’ kernel; the obtained deconvolved kernel ‘minimum’ providing an approximate lower boundary for the original HJ distribution on the sample DNA.

#### Quantitative PCR on ADP1 region

Used for in vivo trioxsalen crosslinking PCR template inhibition assay, performed with Thermo Scientific 2xMAXIMA SYBR green/Fluorescein qPCR Master Mix (K0241) on a Bio-Rad MyiQ Real-Time PCR machine. gDNA dilutions were quantified by QubitBR, and provided at 240pg in 4ul TE pH7.4 to 25ul reactions in triplicates. The trioxsalen untreated sample was also used for a five 10-fold dilution standard series. ADP1 locus primers at 1pMol each: GGTGATGATTGCTCTCTGCC and CGTCACAATTGATCCCTCCC, amplicon 130bp. Cycling: 1x 95°C 10min; 42x [95°C 30sec, 60°C 52sec, 72°C 30sec + RealTimeAcquisition]; 80x [95°C 20sec -0.7°C + MeltCurveAcquisition]. PCR efficiency: 93.5%.

#### Analysis of the reference genome for direct and inverted repeats

Tandem repeats and palindromes were identified on the ASM205788v1 *S.cerevisiae* SK1 reference genome using EMBOSS v.6.6.0.0, allowing for overlapping repeats. For direct repeat identification, etandem was used, searching for direct repeats 16nt to 170nt in length, using a threshold score of 12 (etandem -sequence $FILES -outfile $FILES’.txt’ - minrepeat 16 -maxrepeat 170 -threshold 12). Tandem reports were converted to bedgraph, adding chromosome identifiers, using zero based start coordinates, and using the tandem score (local identity) as coverage. Palindrome identification used palindrome -sequence SK1_ASM205788v1.fasta -outfile palSK1ASMv1 -minpallen 12 -maxpallen 100 -gaplimit 170 -nummismatches 2, (12-100nt length, maximum gap of 170nt, allowing two mismatches). Reports were converted to bedgraph by retaining only coordinates, using zero-based start coordinates, adding chromosome identifiers, and scoring the length of each individually identified inverted repeat element. For both, tandem and palindrome tracks, overlapping repeats regions were cumulatively scored and uniformly binned at 25nt.

### DETAILED HJSEQ WORKFLOW

Detailed procedures and protocols will be made available upon manuscript acceptance.

---

**Table S1.** Yeast Strain Genotypes.

|  |  |  |
| --- | --- | --- |
| FKY4027 | MATa HO zip3Δ::KanMX4<br>-----<br>MATalpha HO zip3Δ::KanMX4 | Fig.S2<br>Fig.5, S5 |
| FKY6386 | MATa ho::LYS2 lys2 ura3 leu2::hisG trp1::hisG<br>-----<br>MATalpha ho::LYS2 lys2 ura3 leu2::hisG trp1::hisG<br>his3::hisG zip1Δ::KanMX4<br>-----<br>his3::hisG zip1Δ::KanMX4 | Fig.S2<br>Fig.5, S5 |
| MJL3538<br>(YJM15383) | MATa lys2 ho::LYS2 ura3Δ(hind3-sma1)<br>-----<br>MATalpha lys2 ho::LYS2 ura3Δ(hind3-sma1)<br>arg4Δ(eco47III-hpa1) his4Δ(Sal1-Cla1)::URA3rev-tel-ARG4<br>-----<br>arg4Δ(eco47III-hpa1) his4Δ(Sal1-Cla1)::URA3rev-tel-ARG4<br>dmc1Δ::LEU2<br>-----<br>dmc1Δ::LEU2 | Fig.2, S2<br>Fig.4 |
| MJL3554 | MATa lys2 ho::LYS2 ura3Δ(hind3-sma1)<br>-----<br>MATalpha lys2 ho::LYS2 ura3Δ(hind3-sma1)<br>arg4Δ(eco47III-hpa1) his4Δ(Sal1-Cla1)::URA3rev-tel-ARG4<br>-----<br>arg4Δ(eco47III-hpa1) his4Δ(Sal1-Cla1)::URA3rev-tel-ARG4 | Fig.S2<br>Fig.S3<br>Fig.4, S4 |
| MJL3555 | MATa lys2 ho::LYS2 ura3Δ(hind3-sma1)<br>-----<br>MATalpha lys2 ho::LYS2 ura3Δ(hind3-sma1)<br>arg4Δ(eco47III-hpa1) his4Δ(Sal1-Cla1)::URA3rev-tel-ARG4<br>-----<br>arg4Δ(eco47III-hpa1) his4Δ(Sal1-Cla1)::URA3rev-tel-ARG4<br>spo11-Y135F-HA3-His6-KanMX4<br>-----<br>spo11-Y135F-HA3-His6-KanMX4 | Fig.S2<br>Fig.S3<br>‡Fig.4&S4 |
| MJL4018 | MATalpha ho::LYS2 lys2 ura3 arg4Δ(eco47III-hpaI)<br>-----<br>MATa ho::LYS2 lys2 ura3Δ(hind3-sma1) arg4Δ(eco47III-hpa1)<br>leu2R::URA3rev-tel-ARG4 ndt80Δ(eco47III-bseR1)::kanMX6 cyh2-z<br>-----<br>leu2-R ndt80Δ(eco47III-bseR1)::kanMX6 CYH2<br>STE50-NatMX-RRP7 his4::URA3rev-tel-ARG4 pac1(62528)-Sph1<br>-----<br>STE50 RRP7 HIS4 | Fig.S3<br>Fig.4, S4 |
| MJL4091 | MATalpha ho::LYS2 lys2 arg4Δ(eco47III-hpaI) STE50 RRP7<br>-----<br>MATa ho::LYS2 lys2 arg4Δ(eco47III-hpaI) STE50-NatMX-RRP7<br>-----<br>spo11-Y135F-HA::URA3 ndt80Δ(eco47III-bseR1)::kanMX6 cyh2-z<br>-----<br>spo11-Y135F-HA::URA3 ndt80Δ(eco47III-bseR1)::kanMX6 cyh2-z<br>-----<br>HIS4 leu2-R::URA3rev-tel-ARG4 ura3<br>-----<br>his4::URA3rev-tel-ARG4 leu2-R ura3 | ‡Fig.S3<br>Fig.4, ‡S4 |
| YJM12947 | MATa ho::LYS2 lys2 ura3 leu2::hisG trp1::hisG<br>-----<br>MATalpha ho::LYS2 lys2 ura3 leu2::hisG trp1::hisG<br>-----<br>fpr1Δ::NatMX4 RPL13A-2xFKBP12::TRP1 REC104-FRB-HA3::KANMX6<br>-----<br>fpr1Δ::NatMX4 RPL13A-2xFKBP12::TRP1 REC104-FRB-HA3::KANMX6<br>-----<br>ndt80::HIS3 KanMX4::pGAL1-Yen1-9A-myc18::URA3<br>-----<br>ndt80::HIS3 KanMX4::pGAL1-Yen1-9A-myc18::URA3<br>-----<br>his3::GPDp-GAL4(484).ER-HIS3<br>-----<br>his3::GPDp-GAL4(484).ER-HIS3 | Fig.4, S4 |
| YJM14569 | MATa ho::LYS2 lys2 ura3 leu2::hisG trp1::hisG<br>-----<br>MATalpha ho::LYS2 lys2 ura3 leu2::hisG trp1::hisG<br>-----<br>fpr1Δ::NatMX4 RPL13A-2xFKBP12::TRP1 REC104-FRB-HA3::KANMX6<br>-----<br>fpr1Δ::NatMX4 RPL13A-2xFKBP12::TRP1 REC104-FRB-HA3::KANMX6<br>-----<br>ndt80::HIS3 KanMX4::pGAL1-Yen1-9A-myc18::URA3<br>-----<br>ndt80::HIS3 KanMX4::pGAL1-Yen1-9A-myc18::URA3<br>-----<br>his3::GPDp-GAL4(484).ER-HIS3 spo11::KITRP1<br>-----<br>his3::GPDp-GAL4(484).ER-HIS3 spo11::KITRP1 | Fig.4, S4 |
| YJM15213 | MATa HO<br>-----<br>MATalpha HO | Fig.1-5<br>Fig.S1-S5 |
| YJM15214 | MATa HO spo11-Y135F-HA3-His6::KanMX4<br>-----<br>MATalpha HO spo11-Y135F-HA3-His6::KanMX4 | Fig.1-5<br>Fig.S1<br>‡Fig.S2<br>Fig.S3<br>‡Fig.S4&S5 |
| YJM15215 | MATa HO ndt80Δ::NatMX4<br>-----<br>MATalpha HO ndt80Δ::NatMX4 | Fig.2, S2<br>Fig.3, S3<br>Fig.4<br>Fig.5, S5 |
| YJM15217 | MATa HO ndt80Δ::NatMX4 spo11-Y135F-HA3-His6::KanMX4<br>-----<br>MATalpha HO ndt80Δ::NatMX4 spo11-Y135F-HA3-His6::KanMX4 | ‡Fig.2, S2<br>‡Fig.3&S3<br>‡Fig.4<br>‡Fig.5&S5 |
| YJM16079 | *MATa* *ho::LYS2 ura3 leu2::hisG trp1::hisG his3::hisG*<br>-----<br>*MATalpha* *ho::LYS2 ura3 leu2::hisG trp1::hisG his3::hisG*<br>-----<br>*ndt80Δ::NatMX4 spo11-Y135F-HA3-His6::KanMX4*<br>-----<br>*ndt80Δ::NatMX4 spo11-Y135F-HA3-His6::KanMX4*<br>-----<br>*sgs1::CLB2p-HA3-SGS1::KanMX6*<br>-----<br>*sgs1::CLB2p-HA3-SGS1::KanMX6* | ‡Fig.S2<br>‡Fig.5&S5 |
| YJM16080 | *MATa* *ho::LYS2 ura3 leu2::hisG trp1::hisG his3::hisG*<br>-----<br>*MATalpha* *ho::LYS2 ura3 leu2::hisG trp1::hisG his3::hisG*<br>-----<br>*ndt80Δ::NatMX4 sgs1::CLB2p-HA3-SGS1::KanMX6*<br>-----<br>*ndt80Δ::NatMX4 sgs1::CLB2p-HA3-SGS1::KanMX6* | Fig.S2<br>Fig.5, S5 |

‡ indicating use for HJSeq as Reference for subtraction only in relevant figure

*S.mikatae* IFO1815
|  |  |  |
| --- | --- | --- |
| FKY6512<br>(SKY2522)<br>(YJM15873) | *MATa* *HO*<br>-----<br>*MATalpha* *HO* | calibration<br>spike-in |

**Table S2.**
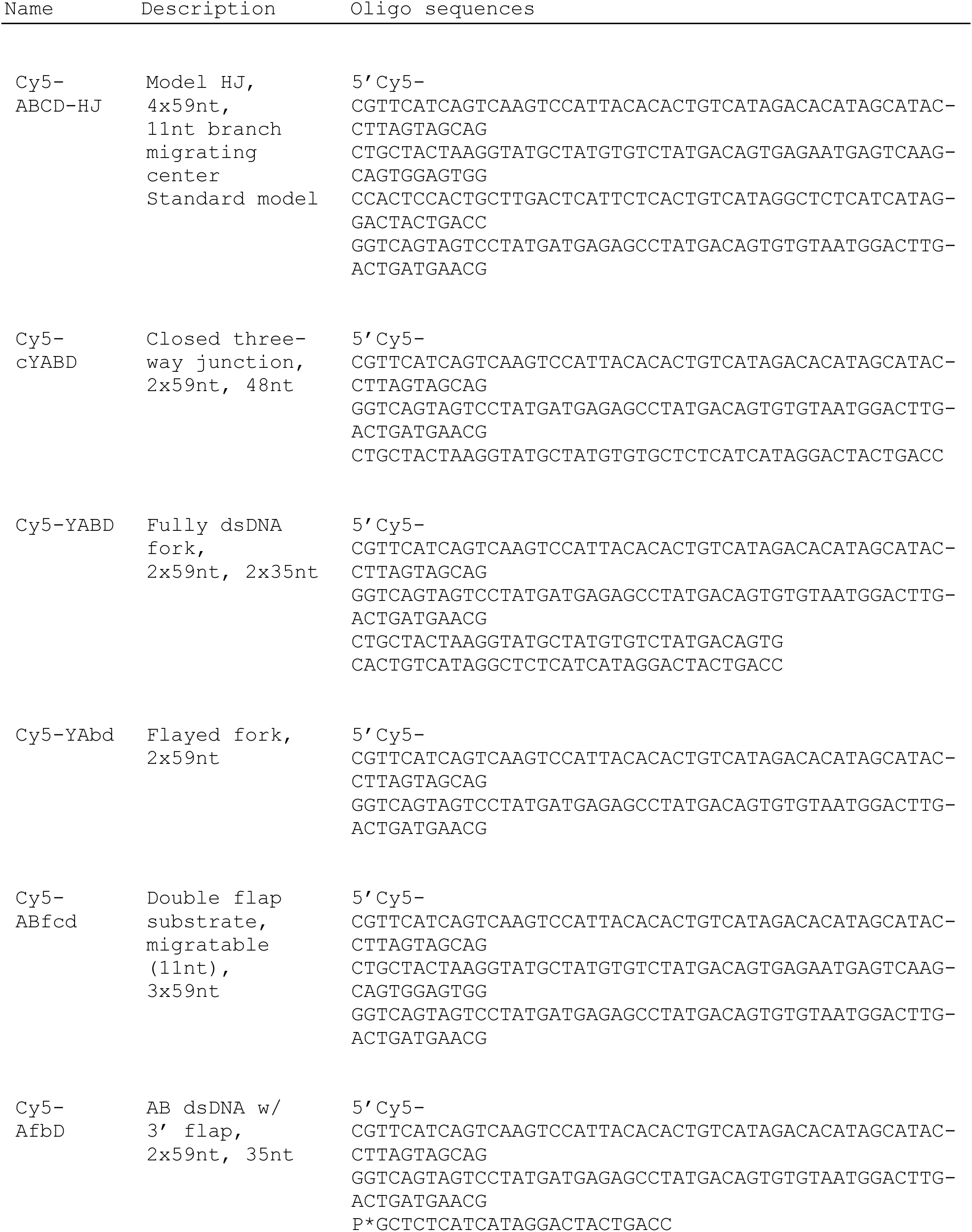
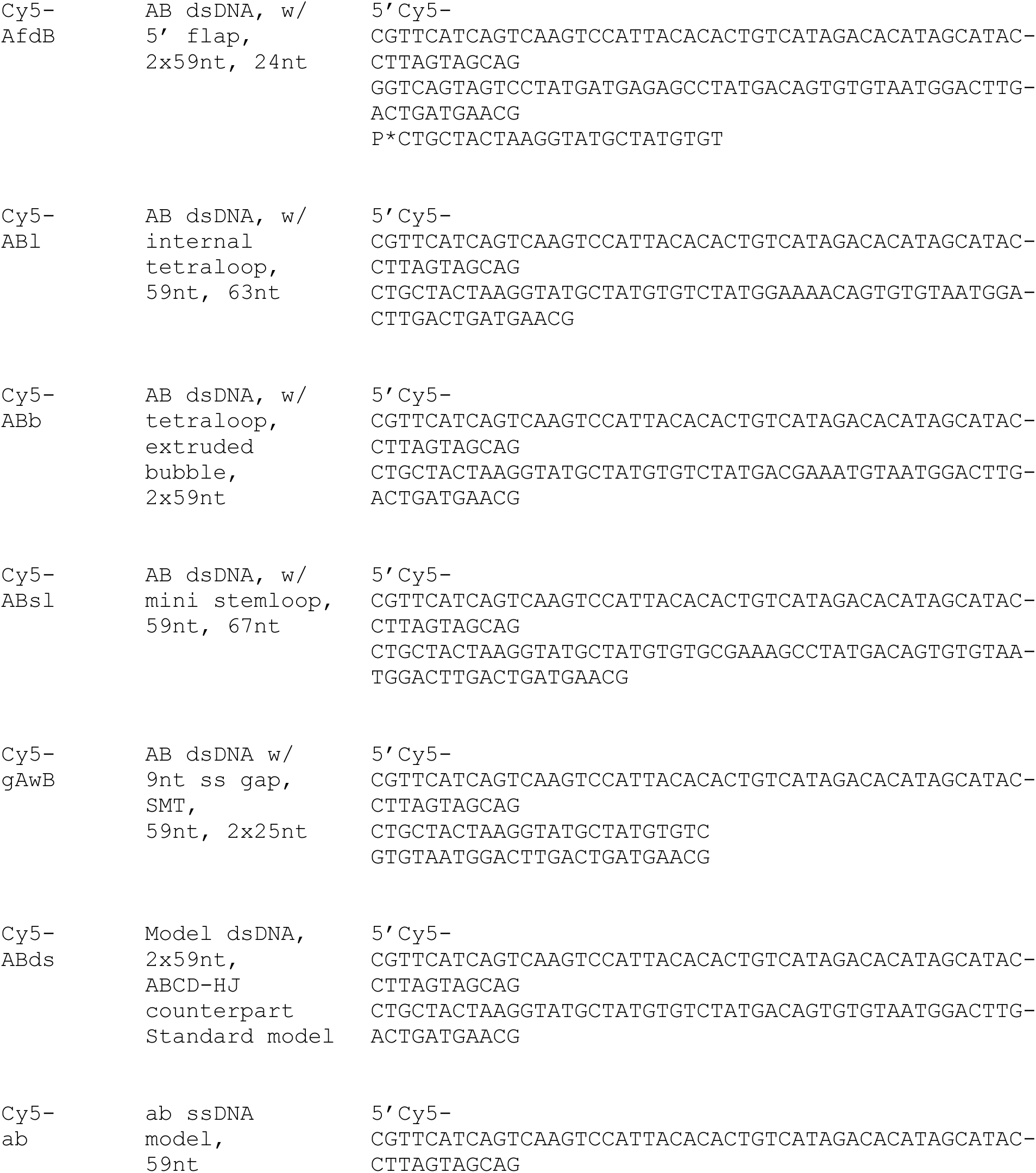
Model substrates.

**Table S3.**
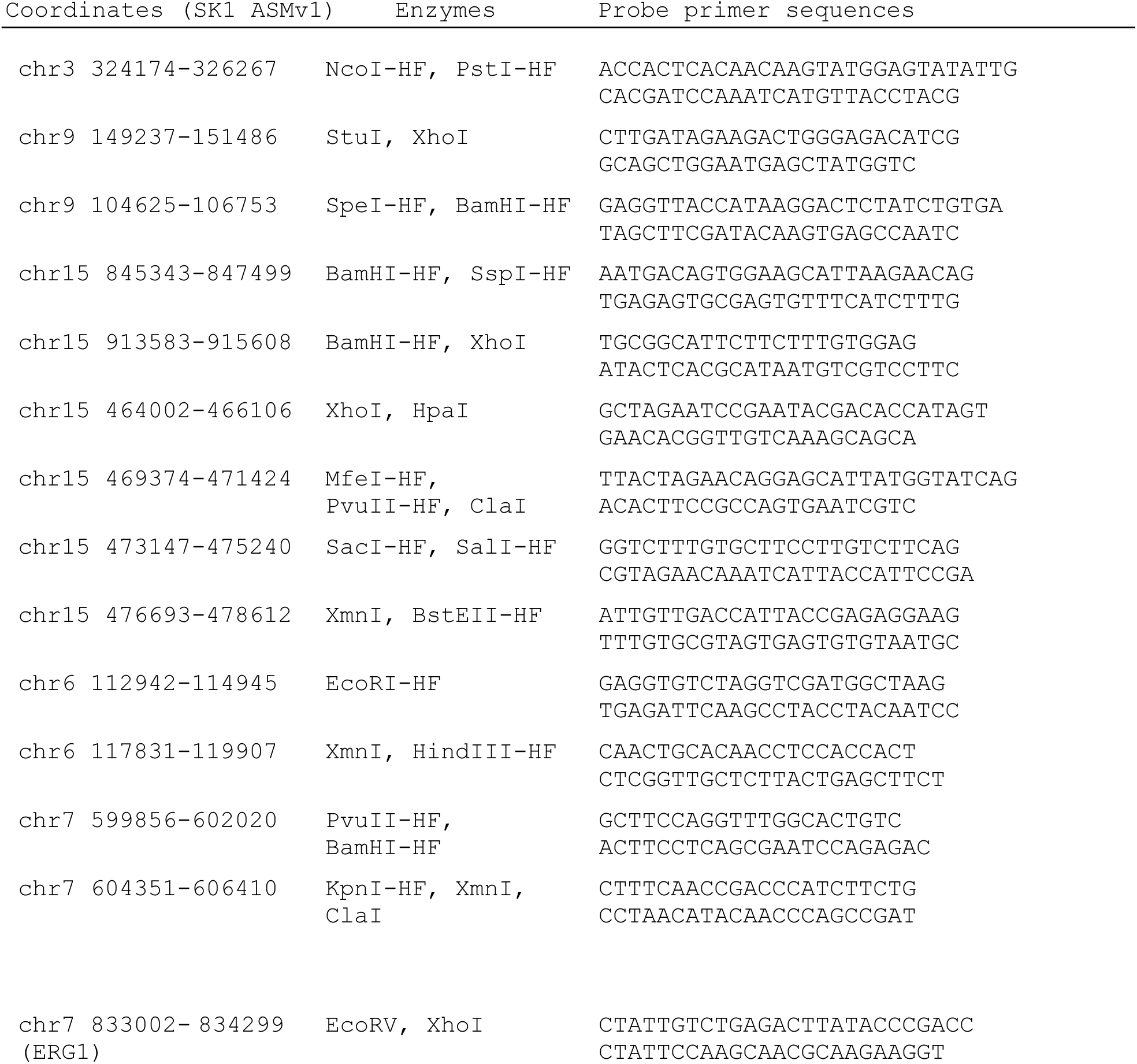
Southern blotting JM assays.

